# High-frame-rate analysis of knee cartilage deformation by spiral dualMRI and relaxation mapping

**DOI:** 10.1101/2022.03.11.483911

**Authors:** Woowon Lee, Emily Y. Miller, Hongtian Zhu, Callan M. Luetkemeyer, Stephanie E. Schneider, Corey P. Neu

## Abstract

**Purpose:** Daily activities including walking impose high frequency cyclic forces on cartilage and repetitive compressive deformation. Analyzing cartilage deformation during walking would provide spatial maps of displacement, strain, and enable viscoelastic characterization, which may serve as imaging biomarkers for early cartilage degeneration when the damage is still reversible. However, the time-dependent biomechanics of cartilage is not well described, and how defects in the joint impact the viscoelastic response is unclear.

**Methods:** We used spiral acquisition with displacement encoding MRI to quantify displacement and strain maps at a high frame rate (40 ms; 25 frames/sec) in tibiofemoral joints. We also employed relaxometry methods (T1, T1ρ, T2, T2*) on the cartilage.

**Results:** Normal and shear strains were concentrated on the tibiofemoral contact area during loading, and the defected joint exhibited larger compressive strains. We also determined a positive correlation between the change of T1ρ in cartilage after cyclic loading and increased compressive strain on the defected joint. Viscoelastic behavior was quantified by the time-dependent displacement, where the damaged joint showed increased creep behavior compared to the intact.

**Conclusions:** Our results indicate that spiral scanning with displacement encoding can quantitatively differentiate the damaged from intact joint using the strain and creep response. The viscoelastic response identified with this methodology could serve as biomarkers to detect defects in joints in vivo and facilitate the early diagnosis of joint diseases such as osteoarthritis.

## INTRODUCTION

Cartilage is a connective tissue which acts to cushion joints from impacts caused by motion and provides lubrication to the joint while transferring load.^1,2^ These functions of cartilage are necessary for daily activities, but are lost with damage of cartilage or diseases including osteoarthritis (OA), which leads to severe disruption in quality of life and chronic pain. OA is a soft tissue degenerative disease found in the knee joint and a severe medical and socioeconomic burden afflicting more than 10% of adults over the age of 60 worldwide.^3^ Current diagnostic tools for OA, including radiography and ultrasound, assess morphological features such as tissue loss and joint space narrowing.^4^ However, these methods of detection are effective only when the damage in cartilage has progressed to an irreversible stage due to the lack of knowledge of cartilage degeneration pathogenesis.

Studies have revealed that the change in mechanical properties of cartilage could be an indicator of OA progression.^5,6^ Thus, diagnostic tools that assess cartilage mechanical properties may provide a promising method of detection. One approach to measure the mechanical properties of cartilage is to indent the surface ex vivo and extract properties related such as stiffness, elastic and shear moduli, and viscoelastic parameters.^7–10^ Studies have shown the elastic and shear moduli of OA cartilage are smaller compared to healthy cartilage.^7,8^ Using atomic force microscopy and optical coherence tomography, the surface roughness and viscoelastic behavior of OA cartilage have been reported.^9,10^ Despite the importance of these findings, the mechanical response of cartilage during rapid motion on intact tissue have not been fully discovered due to the low temporal resolution of the measurements and invasive sample preparation steps involved.

High-speed imaging, referring to enhanced temporal resolution of an image, could greatly benefit OA research in various aspects. First, the images visualize and precisely measure tissue and joint motion in acquired time-course images that can match the actual speed of the physical activity. In addition, high temporal resolution results in less motion artifacts since the acquisition time is shortened, which is especially advantageous when imaging humans in vivo. Lastly, high-speed imaging can extract time-dependent mechanical properties of the tissue including viscoelastic properties, which are critical to investigate in load-bearing tissues such as cartilage.^11,12^

MRI is a non-invasive imaging technique which has a sub-millimeter spatial resolution with high contrast on soft tissue and deep penetration depth. Displacement encoding with stimulated echoes (DENSE) encodes the movement of the tissue occurring during the mixing time (TM) to the MR phase,^13^ and is a particularly useful MRI method for characterization of musculoskeletal tissues.^5,14^ The phase in each encoded direction can later be processed to pixel-level displacement and strain maps. In essence, DENSE MRI maps the mechanical strain inside the tissue with high spatial resolution. This is an innovative approach for joints since mechanical property estimates are mainly based on physical contact, necessitating tissue exposure, whereas DENSE MRI can avoid such procedures making it adaptable to in vivo studies.^14^ Previously, we have demonstrated DENSE MRI on knee joints displaying heterogenous displacement and strain patterns.^15^ In an effort to enhance the temporal resolution, the imaging k-space can be filled by multiple spiral interleaves. Spiral scanning inherently has less motion artifacts since the data is collected from the center of k-space making it optimal for fast-moving tissues such as cardiac muscle.^16–18^

In this paper, we use spiral DENSE MRI to capture displacement and strain maps of bovine knee cartilage with exogenous loading, a method we refer to as displacements under applied loading with MRI (dualMRI).^14^ A custom device applies cyclic loading to knee joints at the speed of a walking cadence (0.5 Hz). This approach successfully outputs multi-frame DENSE MR images with a temporal resolution of 40 ms and provides maps of displacements and strains while a compressive load is applied. We apply this technique to both intact and damaged bovine knee joints in the medial condyle and demonstrate an increase in strain in the tibiofemoral contact area following the application of a defect. We also apply relaxometry methods (T1, T1ρ, T2, T2*) on cartilage to observe their changes after cyclic load and find their correlation to the cartilage mechanical response after damage. Lastly, we demonstrate that spiral DENSE MRI can measure the creep response of cartilage under applied load, which is a potential biomarker for detecting OA at an early stage. Ultimately, we believe this method has a strong potential to be applied in vivo due to fast data acquisition and minimum sample preparation, all while providing properties of articular cartilage related to early degeneration.^19^

## METHODS

### DualMRI: Spiral DENSE MRI with Exogenous Cyclic Loading

DENSE MR images were acquired by a clinical MRI system (3T; Siemens Prisma^fit^) (Figure 1A) with a knee coil (15-channel transmitter/receiver, Siemens) while a customized pneumatic loading device (Figure 1B) cyclically compressed the sample. To mimic the speed of the gait frequency, the cyclic loading was composed of 1 s of compressive loading (Figure 1C) followed by 1 s of unloading. An electric pulse was applied before each loading cycle, triggering the spiral DENSE MRI (Figure 1D)^18^ to start image acquisition. As a result, multi-frame DENSE MR images were collected during the loading time frame (Figure 1C) with a temporal resolution equivalent to 40 ms. The images were collected in the coronal plane and the output data contained magnitude and phase.

**Figure 1.**
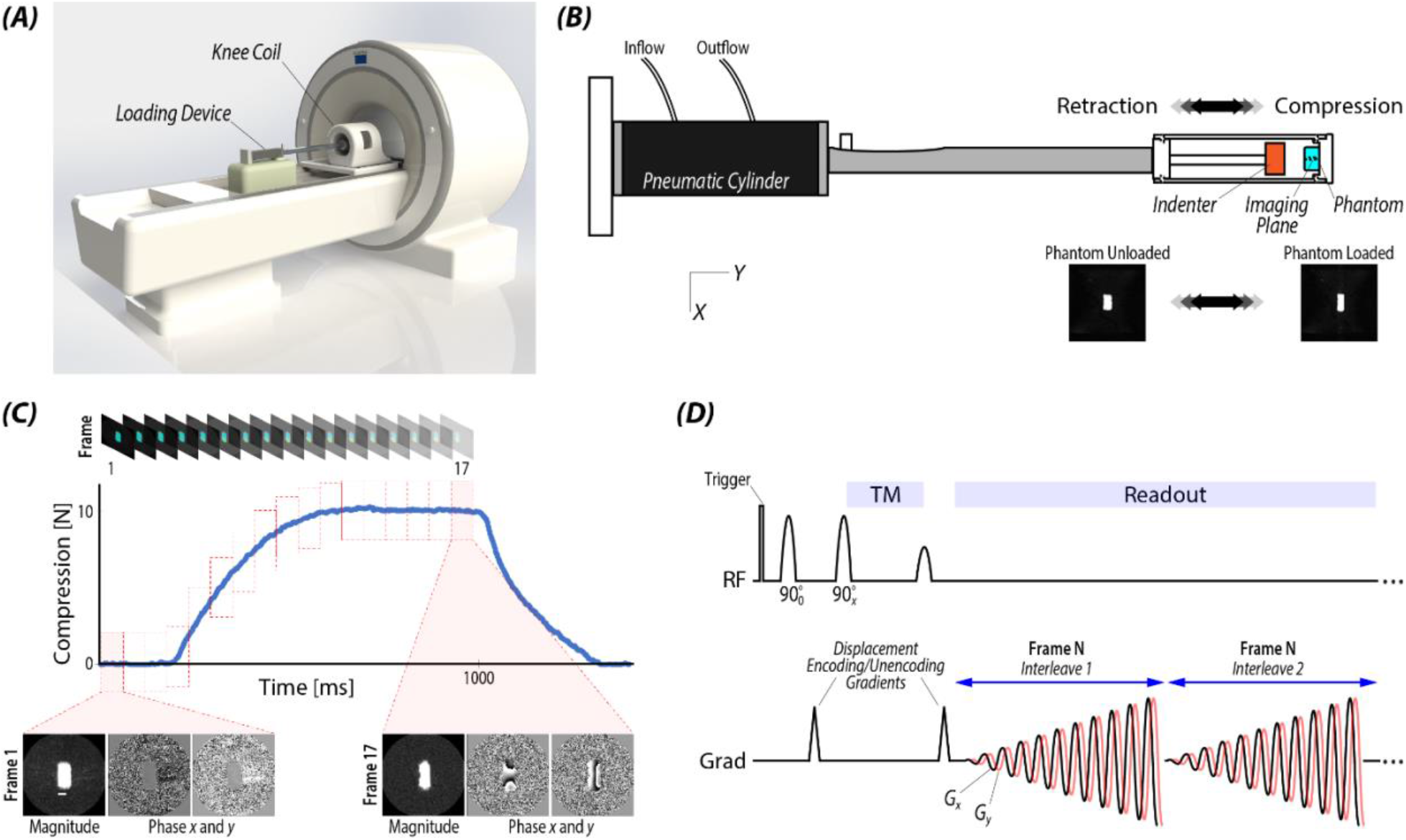
Spiral DENSE MRI acquired high frame rate and displacement-encoded images (40 ms; 25 frames/s) of a phantom sample during mechanical loading. Displacement and strain maps of gel phantom were acquired with exogenous loading, a method we refer to as displacements under applied loading with MRI (dualMRI), at high frame rate (25 frames/s). (A) A 3T Siemens MRI was used to collect DENSE MR images with (B) the pneumatic loading device inside a radiofrequency (RF) knee coil. The phantom loading device had a pneumatic cylinder which controls the cyclic compression and retraction on the phantom by the indenter. (C) Multi-frame DENSE MR images, including magnitude and phase, were obtained throughout the 1 s loading time frame where the first three images had no load due to the physical gap between the indenter and phantom. (D) Deformation during TM was encoded into the phase signal for *x* and *y* where sinusoidal magnetic gradient (Grad) fields were applied to collect data on k-space generating a spiral trajectory. In one loading cycle, two interleaves in k-space were collected per frame. Before TM, gradients were applied in three directions (*x*=0, 2π/3, 4π/3) for displacement encoding to increase SNR and artifact elimination.

Ten spiral interleaves were used to collect data on k-space and two interleaves were acquired for each loading cycle on each frame. Encoding gradients were applied in the *x* and *y* (loading) direction with respect to the coronal plane. Echo time (TE) and relaxation time (TR) were 2.5 and 20 ms, respectively. We used simple three-point displacement encoding and three-point phase cycling for noise reduction.^20^ High signal-to-noise ratio (SNR) double echo steady state (DESS) images were collected before and during constant load for segmentation purpose and to confirm motion of joint tissues to mechanical loading.

### DualMRI Validation on Gel Phantom

A silicone gel phantom made of polydimethylsiloxane (PDMS) (Dow Corning) was prepared using a 1:2 mix ratio of Sylgard 527 and was cut in a 3×3×2 cm^3^ cuboid. The sample was fixed at the end of the loading device for the indenter to apply direct compressive load (10 N) similar to previous design.^15^ For the DENSE MR images, we used a field-of-view (FOV) of 90×90 mm^2^ and total 17 images were collected during the loading time frame. There was a 300 ms time delay between the trigger and image acquisition.

### Displacement and Strain Calculation

Regions of interest (ROIs) for the samples were manually segmented using custom software (MATLAB, 2019b). Displacements for each pixel within the ROIs were determined from phase data as previously described.^15^ Displacements were manually phase unwrapped, smoothed with either a locally weighted scatterplot smoothing (LOWESS) filter (span=10) or Gaussian filter in *x* and *y*. Subsequently, smoothed displacements were used to calculate in-plane Green-Lagrange strains and principal strains.

### Displacement and Strain Precision

Five repeated tests using the loading apparatus were conducted at different combination of parameters that affected the SNR including the number of averages (NA=1, 4, 8), spatial resolution (pixel size=257×257 μm^2^, 360×360 μm^2^, 600×600 μm^2^), and slice thickness (2, 4, 8, 20 mm), while maintaining a fixed FOV (90×90 mm^2^). SNR was measured by first choosing a cartilage and background region in the DENSE MR magnitude image. Subsequently, the mean intensity for both regions was calculated to obtain the ratio. Displacement encoding gradient, ke (0.16, 0.32, 0.64, 0.96 cycles/mm), and smoothing parameters including the number of smoothing cycles (1–100) and types of smoothing (LOWESS, Gaussian) were also controlled to determine the optimal smoothing parameters. To compute precision, 20 evenly spaced points of interest that were consistent across all five scans were selected within the deformed images of the phantom. Experimental precision was defined as the pooled standard deviation at those points of interest across the five repeated tests, and computed as a function of the parameters mentioned above. Precision was calculated for both displacement and strain measurements at the last image frame (17) during the loading cycle.

### Demonstration of Spiral DENSE MRI in Bovine Joints

Seven juvenile bovine joints (1-week-old) were obtained from an abattoir and stored at -80°C. The joints were thawed at 4°C for 48 hs before the experiment and wrapped with phosphate-buffered saline soaked paper towels to prevent dehydration. After the joints were thawed, excessive tissue was removed without damaging the joint capsule, and the tibia and femur was cut on both ends. The tibia and femur were mounted to the sample holders using bone cement (Heraeus) for placement in the pneumatic loading device, similar to a previously published design.^19^ The tibia was mounted on the side facing the indenter. Cyclic loading (100–150 N; 0.5× body weight on three joints and 350–400 N; >1× body weight on four joints) was applied along the superior-inferior axis. Between the trigger and each loading cycle there was a 100 ms time delay to confirm the DENSE MRI data acquisition was initiated before load was applied to the sample. Prior to spiral DENSE MRI acquisition, compressive cyclic loading was applied on the joint for 15 mins to achieve a quasi-steady state response.^21^

The following parameters were used: FOV=125×125 mm^2^, spatial resolution=360×360 μm^2^, ke=0.32 cycles/mm, NA=8, and slice thickness =1.7 mm. We used NA=8 to obtain a high SNR. The total time to acquire one set of DENSE data was 14 mins. Additionally, to simulate cartilage damage, the third sample, which was compressed by 0.5× body weight of load, after MR imaging was opened by single medial parapatellar incision and a critical-sized defect (≈1 cm diameter) was generated on the medial condyle. Subsequently, DENSE MR images were collected on the same (registered) imaging plane with the defect. Displacement fields from all samples went through manual phase unwrapping and LOWESS smoothing for 100 cycles.

### Data Analysis and Statistical Model

The ROIs were divided into total 12 regions based on the type (tibia, femur), direction (lateral, medial) of bone, and if the tibia and femur were in contact or not (contact, non-contact).^22,23^ For the regional analysis of the samples, the average displacement and strain were calculated from the pixels in the region for a total of 12 regions per bovine joint (N=3 intact bovine joints compressed by 0.5× body weight). Standard error of the mean (SEM) was calculated from the average of the three samples. To model the change in displacement and strain at three time points (t=0, 280, 1040 ms), a type II sum of squares ANOVA was run with a linear mixed effects model (nlme package, version 3.1-140) in R (RStudio, version 1.2.1335; R, version 3.6.1). Region was considered a fixed effect and the random effect was each sample. Post-hoc tests were performed with the emmeans package (version 1.4.3.01) using Tukey’s HSD corrections for multiple comparisons. Normality assumptions required for ANOVA were validated before running post-hoc tests. If data did not meet the assumptions of ANOVA, the data was transformed using the cube root transformation and analyses were re-run. To examine the relationship between time and displacement or strain on the intact joints, a linear mixed effects model was used where time was the fixed effect and sample was considered a random effect.

For the intact verses damage dataset, we used a linear mixed model to examine the contact and non-contact regions. Time was treated as a random factor. We then used a type II ANOVA to examine the interaction between contact/non-contact regions with treatment (intact, defect). In the data set, sample is the treatment and contact, non-contact are the divided regions of the knee. At t=1040 ms, 95% confidence intervals (CI) on single bovine joint for the defect and intact comparison were calculated in R. If CI did not overlap, its comparison was considered significant. *Relaxometry: T1, T1ρ, T2, T2*^*\**^

Four types of relaxometry methods (T1, T1ρ, T2, T2^*^) were conducted on all the bovine joints. T1 acquisition parameters were as follows: TE/TR=2.68/15 ms, FOV=80×80 mm^2^, spatial resolution=210×210 μm^2^, slice thickness=1.7 mm, slice number=14, flip angle=5°, NA=2. For T1ρ^24^, the following parameters were used: TE/TR=3/6 ms, spin-lock durations=50, 100, 150, 200 ms, FOV=90×90 mm^2^, spatial resolution=700×700 μm^2^, slice thickness=3.5 mm, slice number=1, flip angle=10°, NA=2. The relaxation map for T1ρ was acquired by fitting the MR intensity images to a monoexponential relaxation model.^25^ Voxels that had coefficient of determination (R^2^) less than 0.66 with respect to the fitting were removed.^26^ T2 mapping parameters were as follows: TEs=13, 26, 39, 52, 65, 78 ms, TR=1290 ms, FOV=80×80 mm^2^, spatial resolution=420×420 μm^2^, slice thickness=1.7 mm, slice number=13, flip angle=180°, NA=2. T2^*^ mapping had the following parameters: Tes=4.46, 11.9, 19.94, 26.98, 34.52, 49.68, 64.84, 80 ms, TR=1270 ms, FOV=80×80 mm^2^, spatial resolution=420×420 μm^2^, slice thickness=2 mm, slice number=11, flip angle=60°, NA=1. All relaxometry methods employed fat suppresion and were measured before and after running spiral DENSE MRI.

T1ρ changes after cyclic loading were measured by calculating the T1ρ difference before and after loading with respect to the initial T1ρ value. To find the correlation between the change of strain and T1ρ after loading, we calculate the strain and T1ρ change for each region as described above and calculated the R^2^ (Excel).

### Measurement of Viscoelastic Response

The LOWESS smoothed displacement fields were further smoothed in time using Gaussian smoothing with a kernel size of 1×1×9 pixels^3^. On the temporally smoothed strain, we fit a natural logarithm curve to the strain-time response during the constant loading time period (480–1040 ms). E_creep_ was calculated from the fitted curve which was defined as the normal strain E_*yy*_ at t=480 ms (E_elastic_; strain only from elasticity) subtracted from E_*yy*_ at t=1040 ms (E_total_; strain reflecting the viscoelastic response). The analysis was conducted on the tibiofemoral contact area (7 regions) for the intact and defected joint (loaded by 0.5× body weight) and for quantitative analysis, we used two-tailed t-test assuming unequal variances.

## RESULTS

### Validation on Deformable Phantoms

The displacements and strains in 2D (Figure 2A) calculated from the DENSE phase images showed a gradual shift over time. During t=0–360 ms (frame 1–10), the displacement maps gradually evolved while during t=400–640 ms (frame 11–17) the displacements did not show any drastic changes. The phantom had a displacement pattern displaying a barrel shape where the center of phantom extended on each side for around 1 mm in *x* and minimum displacement near the surface due to friction. In the loading direction (*y*), the surface nearer to the indenter had high displacement (2 mm) while the fixed surface had minimal displacement (<0.02 mm). E_*yy*_ reached a maximum value close to -0.3 while E_*xx*_ was around 0.1. Both normal strains were concentrated near the center of the phantom. Shear strain (E_*xy*_) showed a more heterogenous pattern: values ranging from -0.1 to 0.1 in the corners. Principal strains 1 and 2 were in the same range with E_*xx*_ and E_*yy*_, respectively since the load was unconfined compression in *y* (Supporting Information Figure S1).

**Figure 2.**
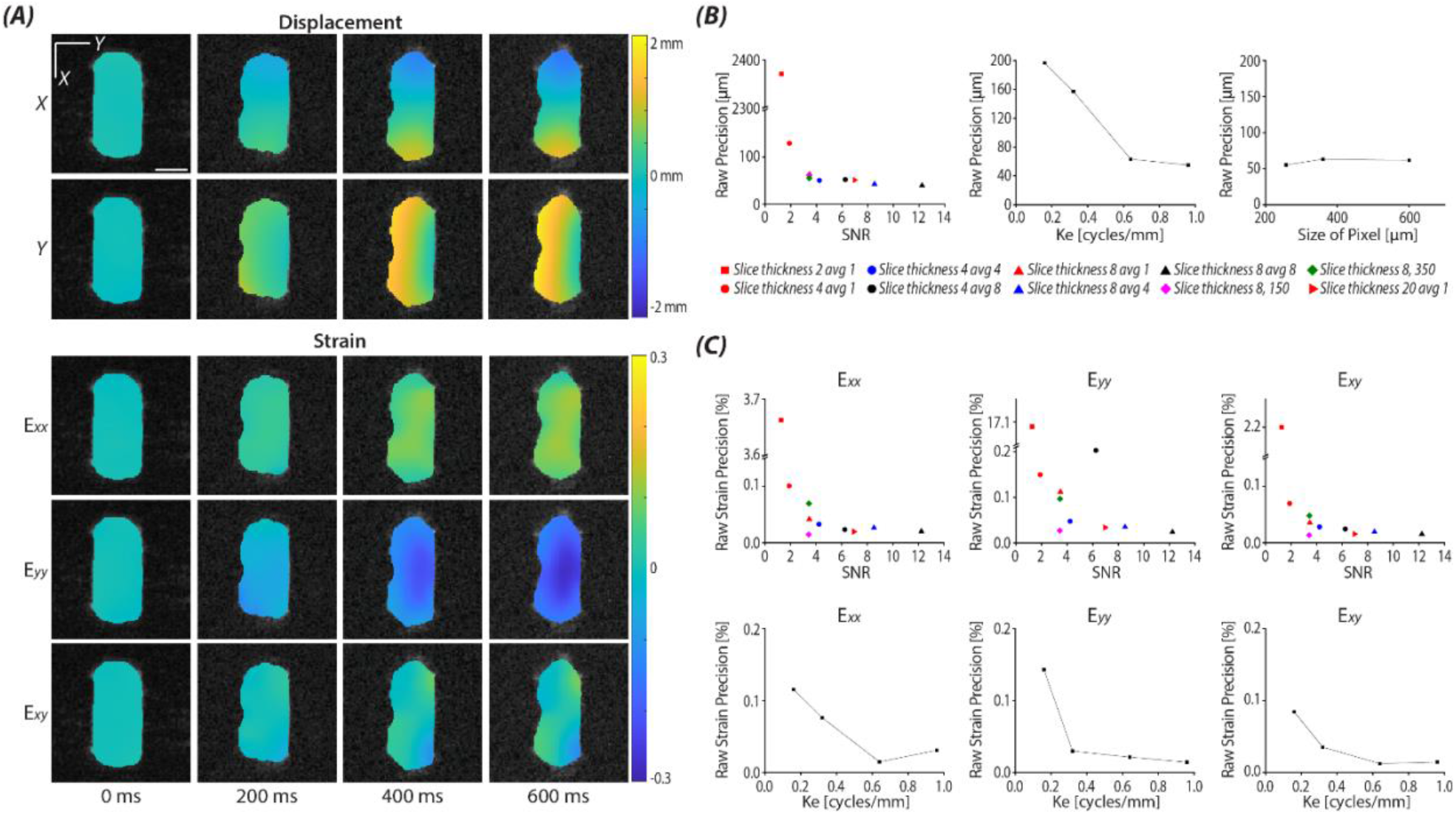
Applying spiral DENSE MRI synced with an external load was validated on a silicone gel phantom. (A) Displacement and strain maps of four representative time points in the loading cycle were obtained by spiral DENSE MRI. The images shown were 4 mm thick, with ke=0.32 cycles/mm, and smoothed 100 times using a LOWESS filter. The displacement and strain maps showed a gradual increase through time. Displacements in *x* and *y* reflected the sample being compressed in *y*. Maximum strain was found on E_*yy*_, which reaches -0.3 while E_*xx*_, and E_*xy*_ were ≈0.1. Scale bar=10 μm. (B) Precision of raw displacement increased with high SNR and ke values, whereas pixel size did not affect the precision. (C) Strain precision also improved with high SNR and ke.

The raw (before smoothing) displacement and strain precision depended strongly on the SNR and ke values (Figure 2B). High SNR improved the raw displacement precision and converged to around 40 μm. High ke values also increased the raw precision, however, this caused more pixels needed for manual phase unwrapping. Pixel size did not affect the raw precision. Both Gaussian and LOWESS smoothing increased the precision by repeating the number of smoothing cycles (Supporting Information Figure S2). Similar precision results were observed from the strain maps (Figure 2C).

### Application on Bovine Joints

Spiral DENSE MRI under cyclic load (0.5× body weight) was able to capture spatially heterogeneous displacement and strain maps on bovine joints altering over time. The loading device used for bovine joints had a similar layout with the phantom loading device with a larger space for the joint sample to fit (Figure 3A). The DESS images displayed the tibia compressing the femur particularly in the tibiofemoral contact area. The time for the applied load to reach the desired value was 400 ms and since there was a 100 ms time delay prior to loading and triggering started at 15 ms, t=485 ms (frame 13) was when the joint was compressed by constant load (Figure 3B). There was minimal displacement in *x* while in *y* (loading direction) the maximum displacement was up to 2 mm in the tibia and 0.5 mm in the femur (Figure 3D; displaying medial condyle labeled in Figure 3C). The displacement maps started to show no rapid change from t=480 ms. The strain maps also had minimal values at t=0 ms and gradually evolved over time. There were heterogeneous patterns in the strain maps during loading while both compression and shear had maximum values in the tibiofemoral contact area (Figure 3D) as seen in previous work.^15^ E_*yy*_ had the highest magnitude approaching -0.35 under load while E_*xx*_ and E_*xy*_ were less than 0.21. Principal strains and max shear strain followed the same trend as E_*xx*_, E_*yy*_, and E_*xy*_ (Supporting Information Figure S3).

**Figure 3.**
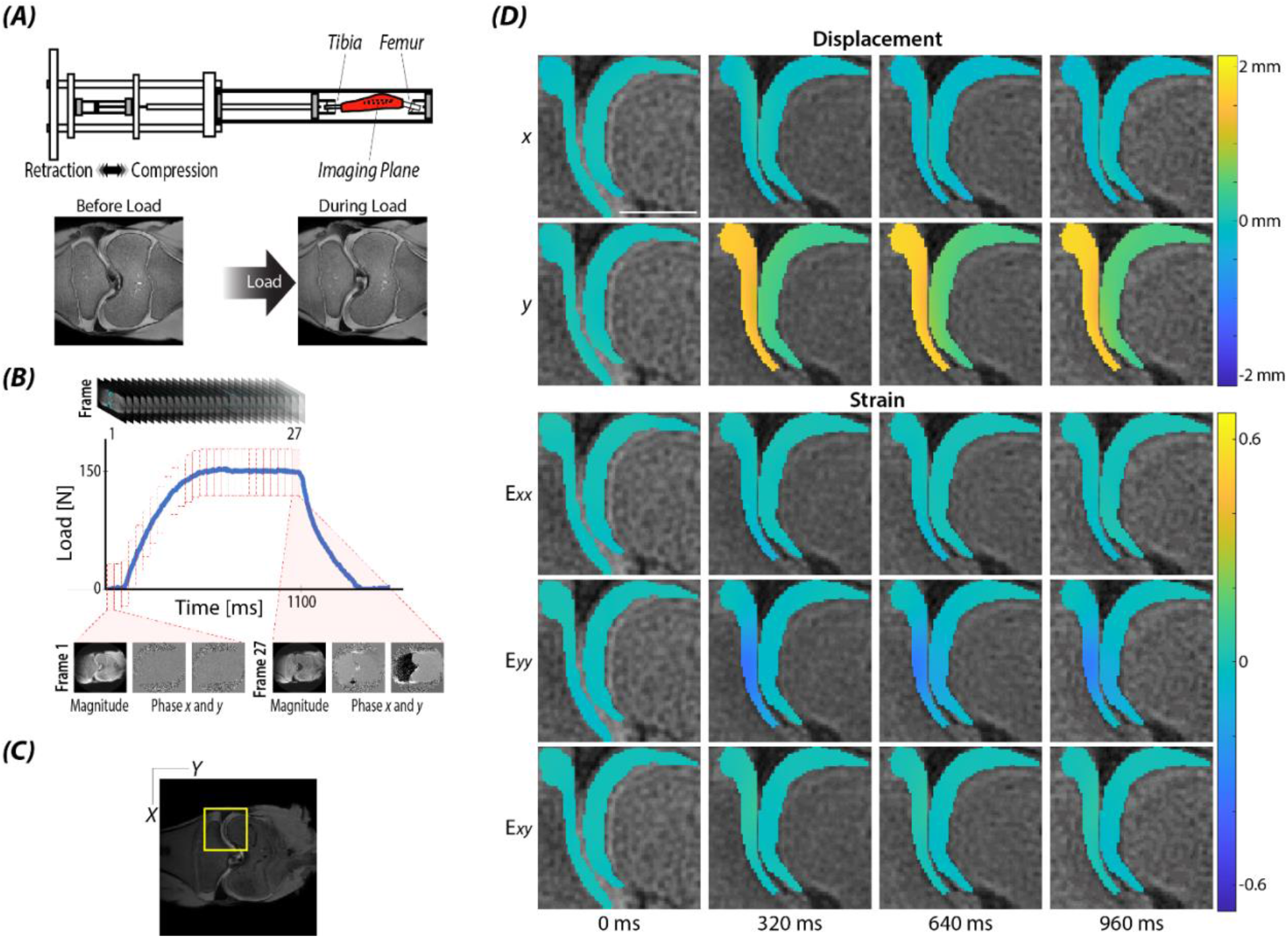
Spiral DENSE MRI demonstrated increased strain during compression of bovine joint samples. (A) The loading device and (B) loading curve were similar to phantom experiments, but with more image frames due to the longer acquisition time (1100 ms). (C) At the medial condyle, (D) the displacement especially in the loading direction *y* led to compressive strains E_*yy*_ in time-course imaging. There was significantly more displacement in the loading direction, *y* compared to *x*. Strain maps were heterogenous and showed maximum values in compressive strain, E_*yy*_, located at the tibiofemoral contact area. Joint samples were loaded by 0.5× body weight. Scale bar=10 μm.

The bovine joints which were loaded more than 1× body weight did not show a gradual change in displacements but rather sudden changes. At t=80–200 ms, there was solely noise in both DENSE magnitude and phase images indicating the sample bent due to the high load and the image slice shifted to a different part of the joint. This is because the joint sample was mounted in a flexed position (flex angle; 20°–30°), higher loads easily caused the sample to bend. As a result, the data was not analyzable for strain calculation and is not shown.

Displacements and strains on joints compressed by 0.5× body weight of load were calculated in localized regions of the ROIs and we found the tibiofemoral contact area had the largest strain during loading for all intact joints. We separated the cartilage into 12 different regions (Figure 4A) to compare the average displacement and strain in each region. One intact sample had roughly 1 mm of displacement in *x* in the tibia region during loading. This affected the average data and was pronounced in the displacement-time curve (Figure 4B). In *y*, the tibial regions had higher displacement values than the femoral regions since the loading indenter was on the tibia. This trend was consistent on all three samples and are shown as distinct lines in the displacement-time curve (Figure 4B). All regions except regions 7 and 8 were statistically significant when comparing t=0 ms verses t=280 ms and t=0 ms verses t=1040 ms in displacement *y*. Both femur (Figure 4C) and tibia contact regions (Figure 4D) displayed a gradual increase in E_*yy*_ with time where tibia contact regions increased greater than femur contact regions reaching -0.12. All regions increased less than 0.03 in E_*xx*_ and E_*xy*_ . Regions 4, 5, and 11 were significant in E_*yy*_ when comparing t=0 ms verses t=280 ms, and t=0 ms verses t=1040 ms. Examining the relation at t=0, 280, and 1040 ms, several regions were significant in displacement *y* and E_*yy*_ (see Supporting Information Table S1-4 for significant region pairs). Principal 1, 2, and max shear strain showed higher magnitudes compared to E_*xx*_, E_*yy*_, and E_*xy*_ but the trend displaying an increase with time was consistent (Supporting Information Figure S4). The displacements, strains, and principal strains for all time points are included in Supporting Information Figures S5–14.

**Figure 4.**
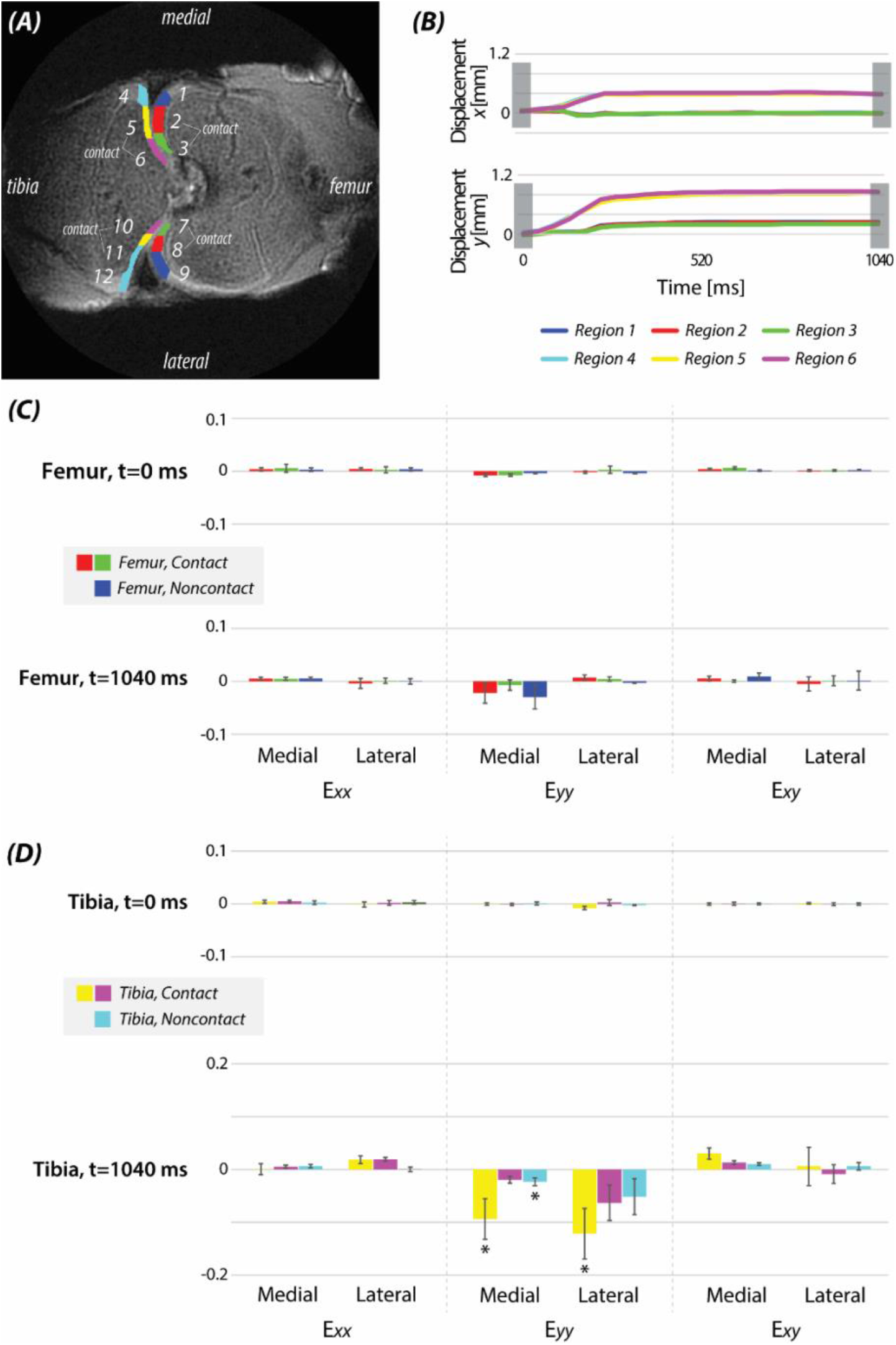
Regional analysis on intact joints showed the increase of strain due to compressive loading (0.5× body weight) was concentrated in tibia contact regions. (A) Articular cartilage was divided into 12 difference regions based on the direction and type of bone. (B) Displacements of the medial compartment showed a gradual increase over time both in *x* and *y*. (C) Normal and shear strains in the femur regions showed an increase at t=1040 ms compared to t=0 ms. (D) Similar with femur regions, all strains in the tibia regions increased at t=1040 ms compared to t=0 ms with greater magnitudes. Statistically significant regions are marked by an asterisk (*). Error bars=SEM (N=3).

The damaged joint showed higher displacements and strains compared to the intact joint. There was more displacement in *x* (Figure 5A) compared to the intact sample (Figure 3D) specially in the tibia (>1 mm) while the displacement in *y* was around 2 mm comparable with the intact sample. Strain maps of the damaged joint displayed heterogeneous patterns where high strains were observed in the tibiofemoral contact area. This trend was more pronounced in E_*yy*_ (labeled with an arrow in Figure 5A). The maximum magnitude values for E_*xx*_, E_*yy*_, and E_*xy*_ were 0.4315, -0.4944, and 0.4623, respectively. Principal 1, 2, and max shear strain showed a comparable trend with E_*xx*_, E_*yy*_, and E_*xy*_, respectively (Supporting Information Figure S15). The ROIs were segmented into 11 regions for regional analysis; the absence of region 2 was due to the defect in the medial condyle (Figure 5B). After comparing the average strains for the contact and non-contact regions with the intact joint, we found the damaged joint had significantly higher E_*yy*_ values in the tibiofemoral contact regions (*P*<0.05) but not in the non-contact regions (*P*>0.05). At t=1040 ms, the damaged sample expressed higher E_*yy*_ in regions 6, 7, 10, and 11 (Figure 5C). A comparison of the intact versus damaged sample in displacements and strains over time for all regions is shown in Supporting Information Figures S16–23.

**Figure 5.**
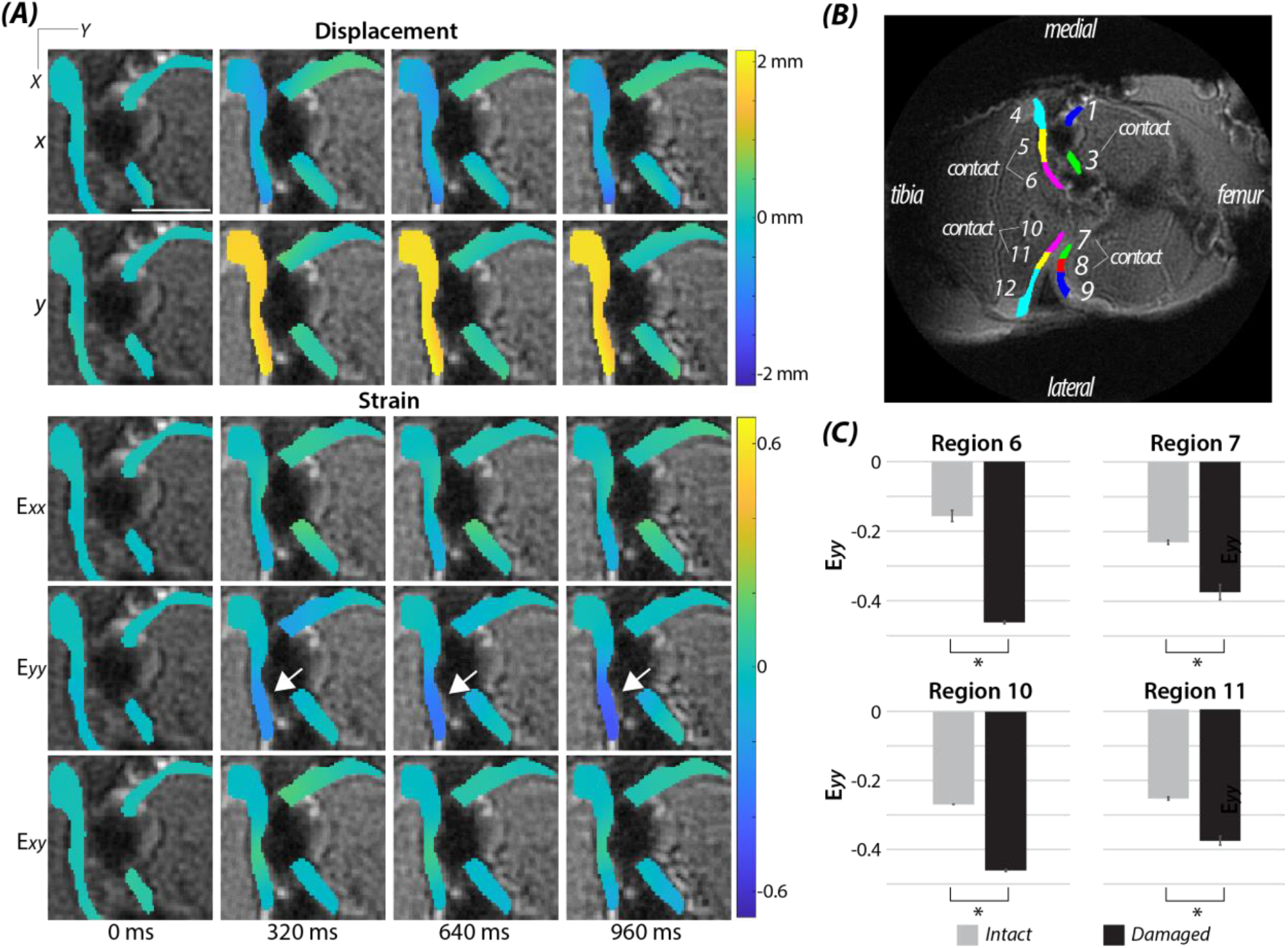
Spiral DENSE MRI showed an increase in strain on the defected joint by compressive load (0.5× body weight) compared to the intact joint. (A) Displacement and strain values gradually evolved over time where E_*yy*_ at t=960 ms showed higher intensities (−0.462) compared to the intact sample (−0.156). Scale bar=10 μm. Strain particularly showed concentration near the defect, marked by the white arrow. (B) The ROIs of the defected sample was divided into 11 regions, excluding region 2 in the intact joint due to the defect. There was a significant difference in (C) E_*yy*_ at the contact regions on t=1040 ms including regions 6, 7, 10, and 1 indicated by an asterisk (*). Error bars=95% CI and N=1.

### Relaxometry

Among the four relaxometry methods applied on cartilage (Figure 6A), T1ρ showed a noticeable increase after loading when the experimental condition was detrimental to the joint. The average values (Figure 6B) for all intact samples were 815 ms, 82 ms, 77 ms, and 36 ms for T1, T1ρ, T2, and T2*, respectively. The average values for each relaxometry method were calculated by measuring the mean value for each pixel within the ROIs. T1, T2, and T2* showed minimal change (<14%) after multiple (>600) cyclic loading whereas T1ρ increased more than 30% when the load was more than 1× body weight or the sample was damaged (Figure 6B). Conversely, there was no significant increase on samples loaded with 0.5× body weight. By plotting the relationship with the change of T1ρ with respect to the change of strain due to damage (Figure 6C), E_*yy*_ had the strongest correlation (R^2^=0.6999) followed by E_*xy*_ (R^2^=0.3987) and E_*xx*_ (R^2^=0.3627).

**Figure 6.**
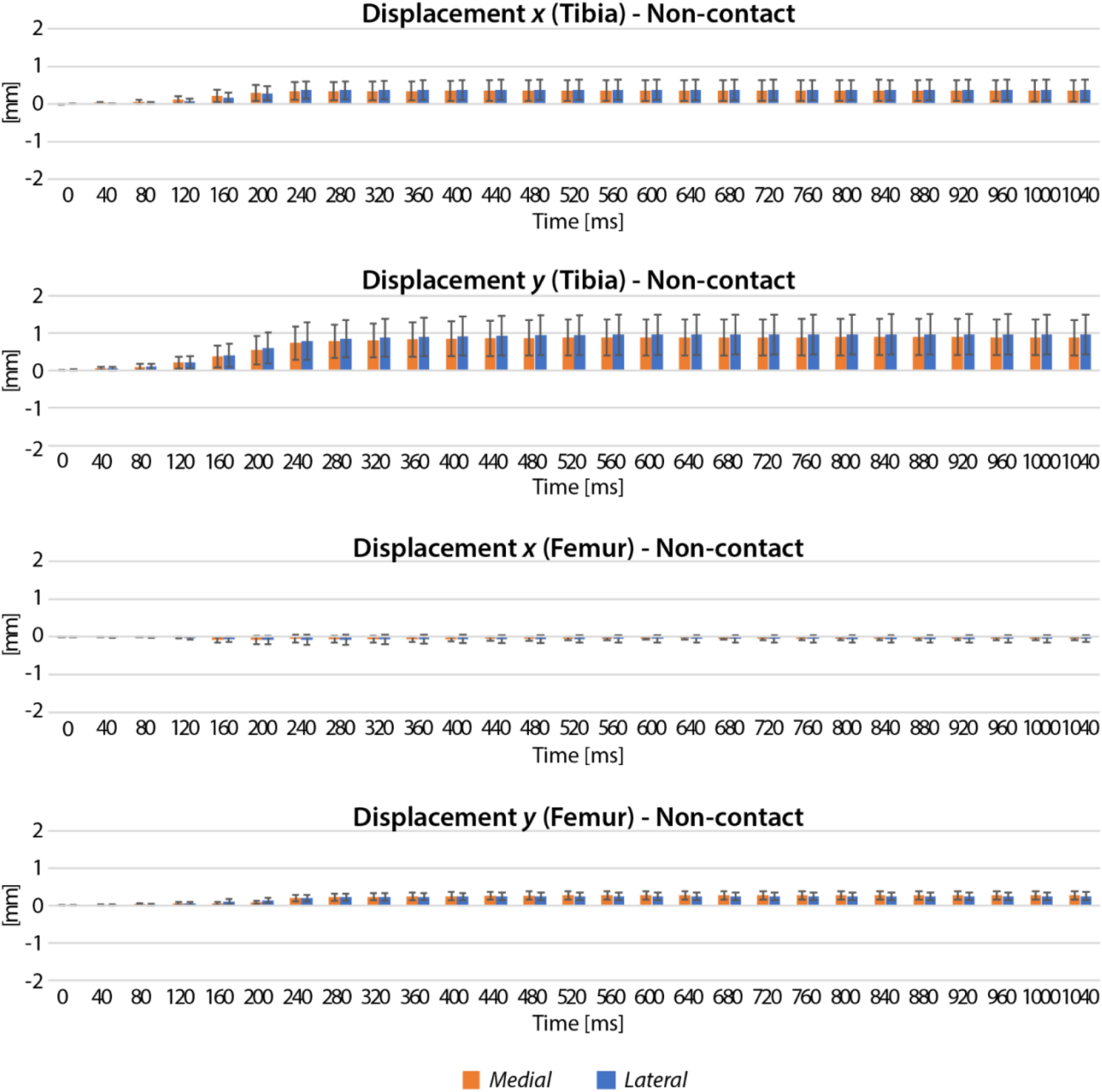
Relaxometry methods on the bovine joint cartilage outputted a pixel-scale map while T1ρ measurements reflected the condition of the joint after cyclic load. (A) T1, T1ρ, T2, and T2* showed a heterogeneous distribution on the cartilage. (B) The average value of the pixels in the selected ROIs on cartilage were plotted. Error bars=SEM. N=4-7 bovine joints. T1ρ increased when more than 1× body weight of load was applied for cyclic loading or when the sample was defected. (C) The increase of E_*yy*_ after damage in each region (labeled in plot) was strongly correlated with the increase of T1ρ. The change of E_*xx*_ and E_*xy*_ after damage showed a weak correlation with the T1ρ increase.

### Viscoelastic Creep Response

The creep behavior of cartilage under compressive load was more pronounced in the damaged compared to the intact joint. Bovine cartilage is a viscoelastic material; thus, during constant load, the strain will exhibit a creep response (Figure 7A). With the improved temporal resolution of spiral DENSE MRI, we were able to quantify the creep behavior (Figure 7B) and compare the E_creep_ values with the intact and damaged sample in the contact regions. The intact sample displayed an average E_creep_ of -0.019 while the damaged sample averaged -0.057 (Figure 7C). However, the difference was not significant (*P*>0.05), as a result of the wide range of whiskers demonstrated in Figure 7C.

**Figure 7.**
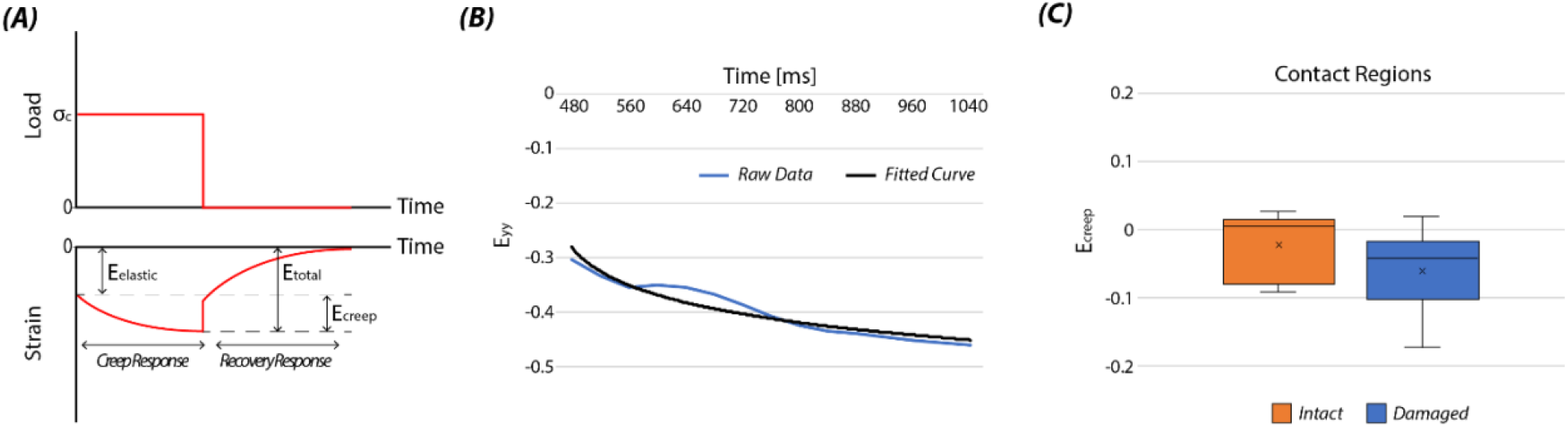
Spiral DENSE MRI enabled to quantify the viscoelastic response of cartilage under compressive load and showed an increased creep response in the damaged joint. (A) E_creep_ was defined as the amount of increase of strain during constant load (0.5× body weight) and was calculated by subtracting E_elastic_ from E_total_. (B) E_*yy*_ strain from region 6 of the defected sample clearly showed a creep response over time and a logarithm curve was fitted. By comparing the contact regions of the intact and damaged sample, (C) the damaged sample had greater E_creep_ values than intact, where the average was -0.019 and -0.057 for intact and damaged, respectively. Statistically, there was no significant difference (paired, two-tail t-test, *P*>0.05) and the mean and median is indicated as × and – in the box plot. Whiskers represents the range of the data points.

## DISCUSSION

This paper demonstrates that spiral DENSE MRI can be applied with external cyclic load and can yield high temporal resolution displacement maps. We validated this approach on a phantom and subsequently applied the method on intact and damaged bovine knee joints with relaxometry mapping. This approach successfully quantified the mechanical strain and viscoelastic response of cartilage under compression utilizing the time-course displacment maps while the loading frequency was comparable with walking cadence. The biomechancial analysis and relaxometry mapping reveals a significant difference in intact and damaged cartilage indicating their possible use as prospective biomarkers for detecting early cartilage degeneration.

The multi-frame images we obtained by spiral DENSE MRI displayed gradual change of displacements and strains with time. Both the phantom and bovine joints showed pixel-specific displacement and strain maps reflecting the compressive force (0.5× body weight on joints) with a time span of 1 s. The phantom correlated with visual observation during loading while the bovine joints showed spatially heoterogenous strain maps, where highly concentrated strains were observed in the tibiofemoral contact area. The non-contact regions of the articular cartilage had lower strain which could be due to the meniscus mediating the load.^27^ The raw displacement and strain precision measured on the phantom improved with high SNR reaching around 40 μm. This was substantially smaller than the lowest pixel size (257×257 μm^2^) and is comparable with other DENSE scanning methods used in our previous work.^15^ Pixel size did not affect the precison since at slice thickness=8 mm, the SNR was constant regardless of the pixel size. The setting we chose for bovine joints was based on matching the optimal precision results from the phantom and choosing a spatial resolution that can capture features in bovine joints. To select an optimal ke, one should consider the absolute movement of the tissue caused by loading since large ke values can cover only small displacements and large displacements will result in more phase unwrapping. Small ke values have a large displacement coverage but have low precision. We chose ke=0.32 after considering both factors.

Spiral DENSE MRI also was effective at distinguishing an intact from damaged joint. The strain values significantly increased in most tibiofemoral contact regions after damage (Figure 5C). Region 6 located adjacent to the defect area showed an increase in strain, specifically in E_*yy*_. This pattern can be interpreted as rim stress concentration around the defect area which have been reported in other studies.^28–30^ In contrast, the strain in region 5 did not increase since the femur facing region 5 was empty, resulting in less tissue compressing the opposing region. E_*yy*_ in the lateral condyle contact regions also increased. As previously mentioned, the displacement in *x* increased significantly after the defect was generated (Figure 5A). This could be due to the medial condyle area contacting the tibia had been eliminated and resulted in an adjustment of load distribution along the articular cartilage. Thus, the modified load affects not only the defected compartment (medial) but also the non-defected compartment (lateral). Overall, the strain analysis on localized regions indicated compressive load was more detrimental to defected cartilage suggesting spiral DENSE MRI can be used to detect OA patients.

Relaxometry analysis was added due to the bio-chemical content potentially affecting the mechanical strain; water content is related to T1 and T2, collagen alignment is connected to T2, and proteoglycan (PG) is linked to T1ρ and T2*.^31,32^ All T1, T1ρ, and T2* values (Figure 6B) were in range of published results on knee cartilage using a 3T scanner,^33–35^ while T2 values were higher than published literature.^33,34^ This could be due to the young age (juvenile) affecting the results where studies show T2 decreases with age.^36^ As previously mentioned, we see an increase (>30%) in T1ρ after cyclic loading when the joint was compressed by large loads (>1× body weight) or when the joint had defects. This supports the inverse relationship of T1ρ with the PG content^37^ as studies have shown the correlation of T1ρ increase with the OA severity due to the loss of PG in extracellular matrix.^38^ We also found correlations of T1ρ change with the strain increase after cartilage damage indicating T1ρ itself contains pivotal information like the PG content but also is diffusely connected to the mechanical properties of the tissue. Some studies show contradicting results of T1ρ which decreased after physical activities.^39,40^ However, these results were based on in vivo subjects which the joints have anchored collagen fibers in the subchondral bone and more amount of synovial fluid.^26^ This leads to dissimilar fluid flow impacting the relative PG content after motion compared to an ex vivo bovine joint.

Lastly, we showed in addition to the mechanical strain, viscoelastic responses could also be extracted by spiral DENSE MRI. Viscoelasticity is an important property of biological tissues and studies have shown its correlation with tissue degradation and early OA progression.^11,12^ However, experimental studies face technical challenges to obtain viscoelastic responses of tissue, especially in intact or in situ joints. This is because viscoelastic measurements need data acquisition with time which may require special methods not suitable for rapid physiolocal movements or in a clinical setting. On the other hand, spiral DENSE MRI captures high temporal resolution images that can measure not only creep behavoir but also other viscoelastic responses of the tissue including time-dependent deformation^10,41^ and relaxation time^42,43^. Such measurements are connected with cartilage damage and PG content.^10,41–43^ Considering the fact that spiral DENSE MRI has a high penetration depth, it becomes an optimal method to measure viscoelastic behavior while easily tranferable to a clinical setting.

There are few limitations in this study which can be further explored. First, there was a small number of joints used for analysis. Also, this method has been restricted to 2D and more complex motions would involve 3D displacement analysis which result in more imaging and processing time. To further analyze the difference between intact and damaged joints, multiple slices around the defect with increased number of samples applicable for an improved statistical model are necessary. In addition, calculating the displacements and strains was time consuming and labor intensive particually when drawing the ROIs and conducting manual phase unwrapping. This was due to the large temporal data and extremely thin ROIs of cartilage making a few pixels highly affecting the strain calculation resulting in abrupt changes of strain with time and ultimately causing positive E_creep_ values. Advanced software tools such as automatic segmentation^44^ and motion tracking^45^ which can maintain the shape of ROIs over time should be considered. To future explore this work, one can apply this approach to in vivo humans while advanced techniques such as compressed sensing can be utilized to shorten the total imaging time.

## CONCLUSION

In summary we present the versatility of spiral DENSE MRI for obtaining the mechanical response of bovine knee cartilage under cyclic compressive load. Spiral scanning enhances the temporal resolution (on the time scale of 40 ms) outputting multi-frame displacement and strain maps during cyclic load with the speed of walking cadence. The displacement and strain analysis shows an increase between intact and damaged joint. Additionally, the ability to successfully extract multi-frame, pixel-level strain maps of cartilage provides a unique ability to quantify the viscoelastic phenomena. We also find T1ρ increases on extreme loading conditions and find the change of T1ρ is positively correlated with the increase of normal strain after damage occurs. In essence, spiral DENSE MRI with relaxometry provides pivotal information of the cartilage’s condition and future studies could include human subjects with known pathological conditions following knee damage, with a goal of predicting early progression of OA.

## ACKNOWLEDGEMENTS

The authors would like to acknowledge funding from NIH R01 AR063712-08. The authors are thankful to Teryn S. Wilkes for operating the MRI at Intermountain Neuroimaging Consortium located at University of Colorado, Boulder, and Nancy Emery for statistical consultation.

## AUTHOR CONTRIBUTIONS

Conceptualization, W.L., C.P.N.; Methodology, W.L., E.Y.M., C.M.L; Device Manufacturing, E.Y.M.; Statistical Analysis, W.L., S.E.S.; Writing – Original Draft, W.L., C.P.N., Writing – Review & Editing, All Authors; Funding Acquisition, C.P.N.

## SUPPORTING INFORMATION

**Table S1.**
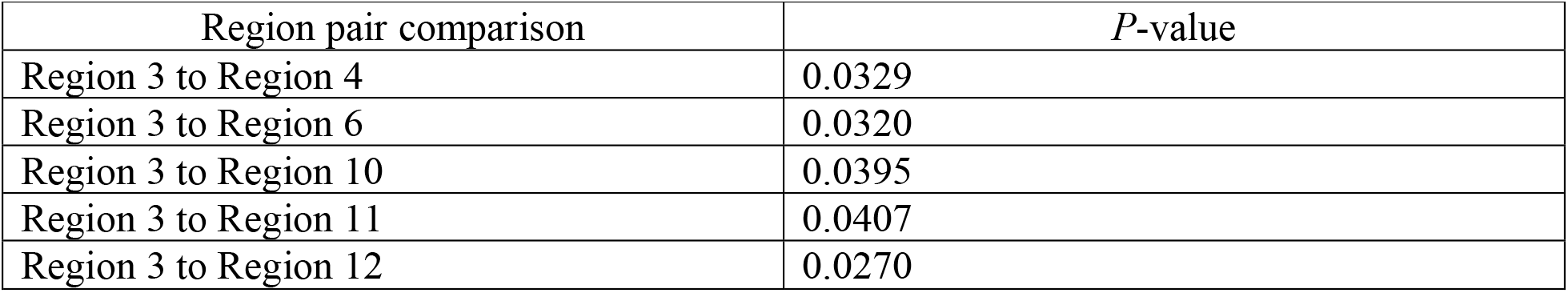
At t=280 ms, displacement *y*.

**Table S2.** At t=280 ms, E_*yy*_.

| Region pair comparison | *P*-value |
| --- | --- |
| Region 2 to Region 5 | 0.0437 |
| Region 3 to Region 5 | 0.0120 |
| Region 4 to Region 5 | 0.0472 |
| Region 5 to Region 9 | 0.0247 |

**Table S3.** At t=1040 ms, displacement *y*.

| Region pair comparison | *P*-value |
| --- | --- |
| Region 3 to Region 12 | 0.04780 |
| Region 6 to Region 7 | 0.0455 |
| Region 7 to Region 12 | 0.0393 |

**Table S4.** At t=1040 ms, E_*yy*_.

| Region pair comparison | *P*-value |
| --- | --- |
| Region 1 to Region 11 | 0.0440 |
| Region 3 to Region 11 | 0.0445 |
| Region 9 to Region 11 | 0.0233 |

**Figure S1.**
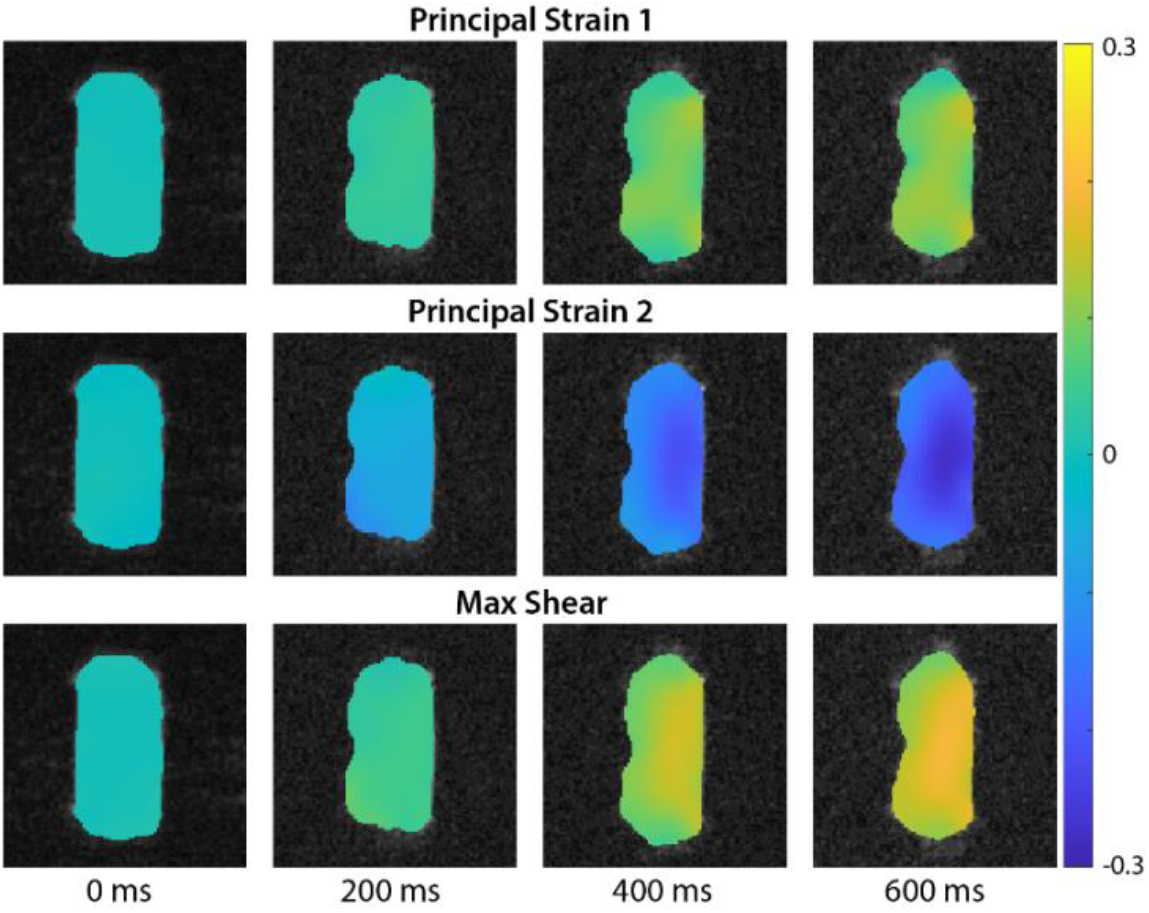
Principal strain 1, 2, and max shear strain obtained by spiral DENSE MRI on a phantom evolved over time. Four representative time points distinctly showed the gradual change of the strains which were similar with E_*xx*_ and E_*yy*_ for principal strain 1 and 2 while max shear strain values reached 0.2.

**Figure S2.**
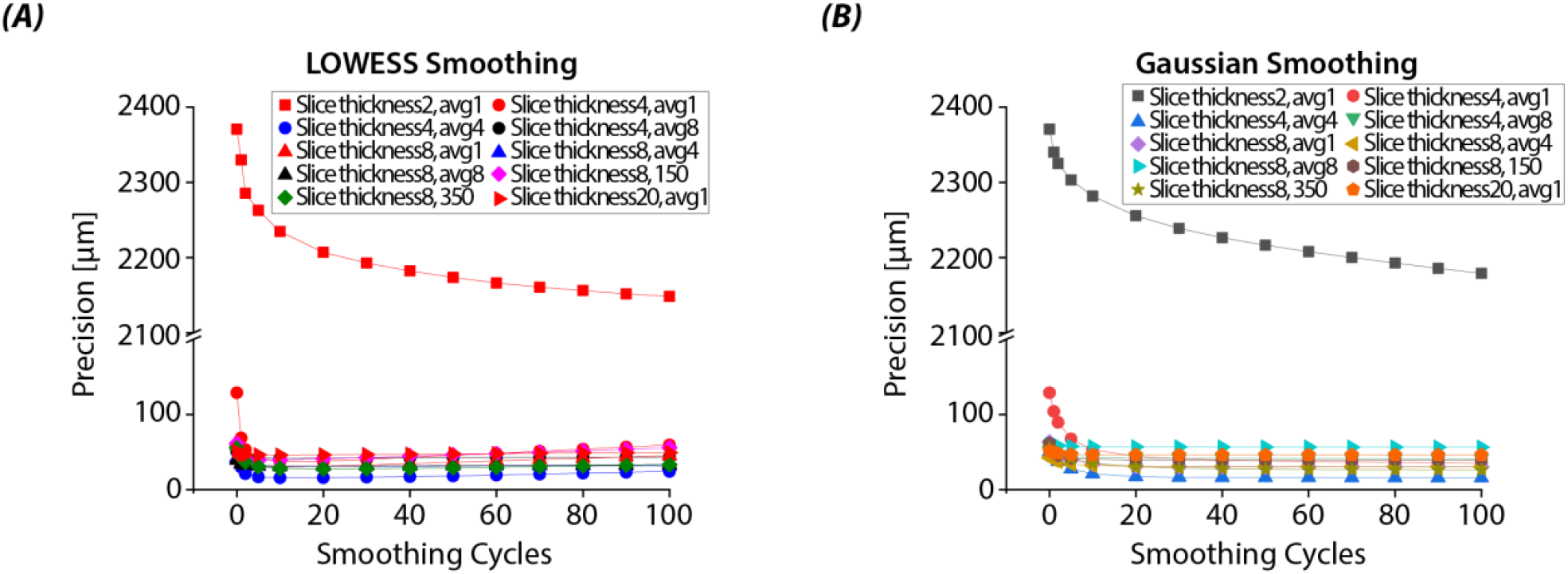
Displacement precision improved over repeated smoothing cycles. Both (A) LOWESS and (B) Gaussian smoothing showed a significant increase of precision which converges to ≈40 μm with repeated number of cycles.

**Figure S3.**
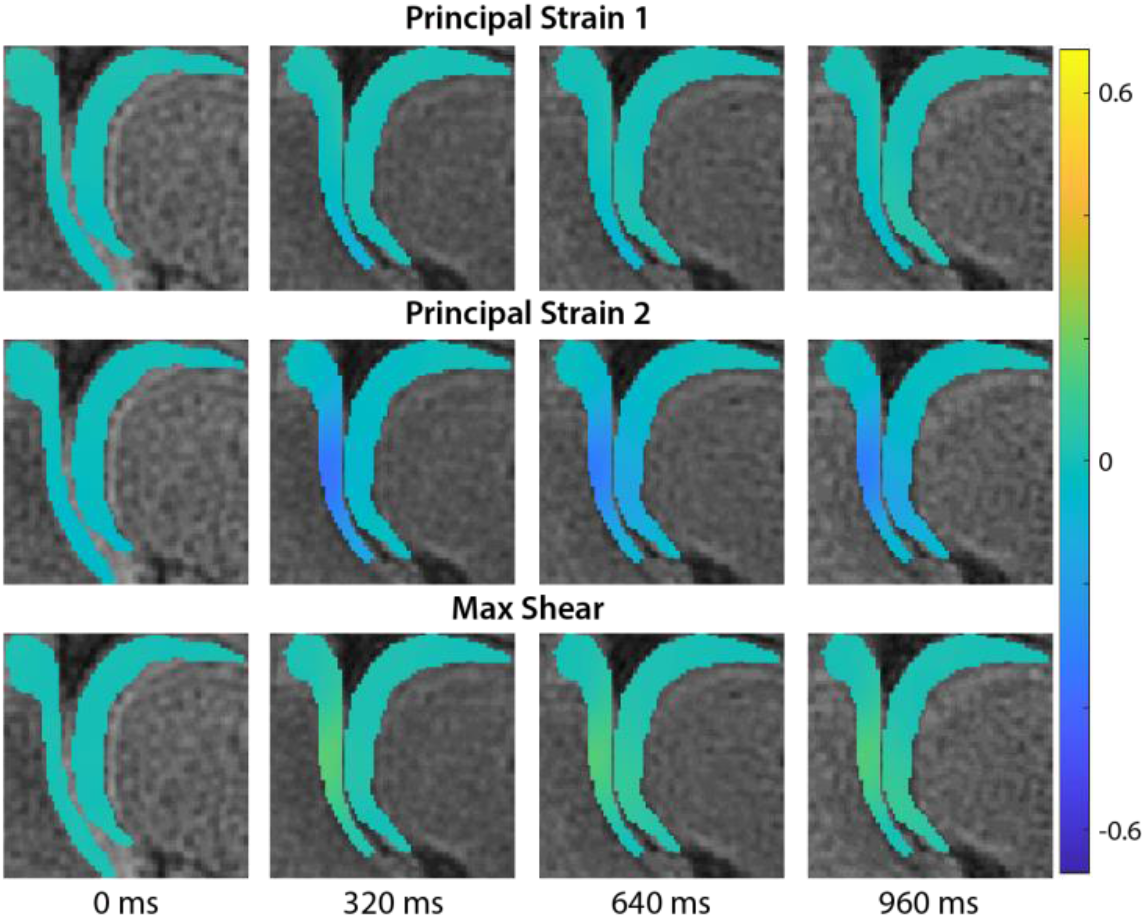
Principal strain 1, 2, and max shear strain obtained by spiral DENSE MRI on an intact bovine joint evolved over time. Strain patterns showed a similar trend with E_*xx*_, E_*yy*_, and E_*xy*_.

**Figure S4.**
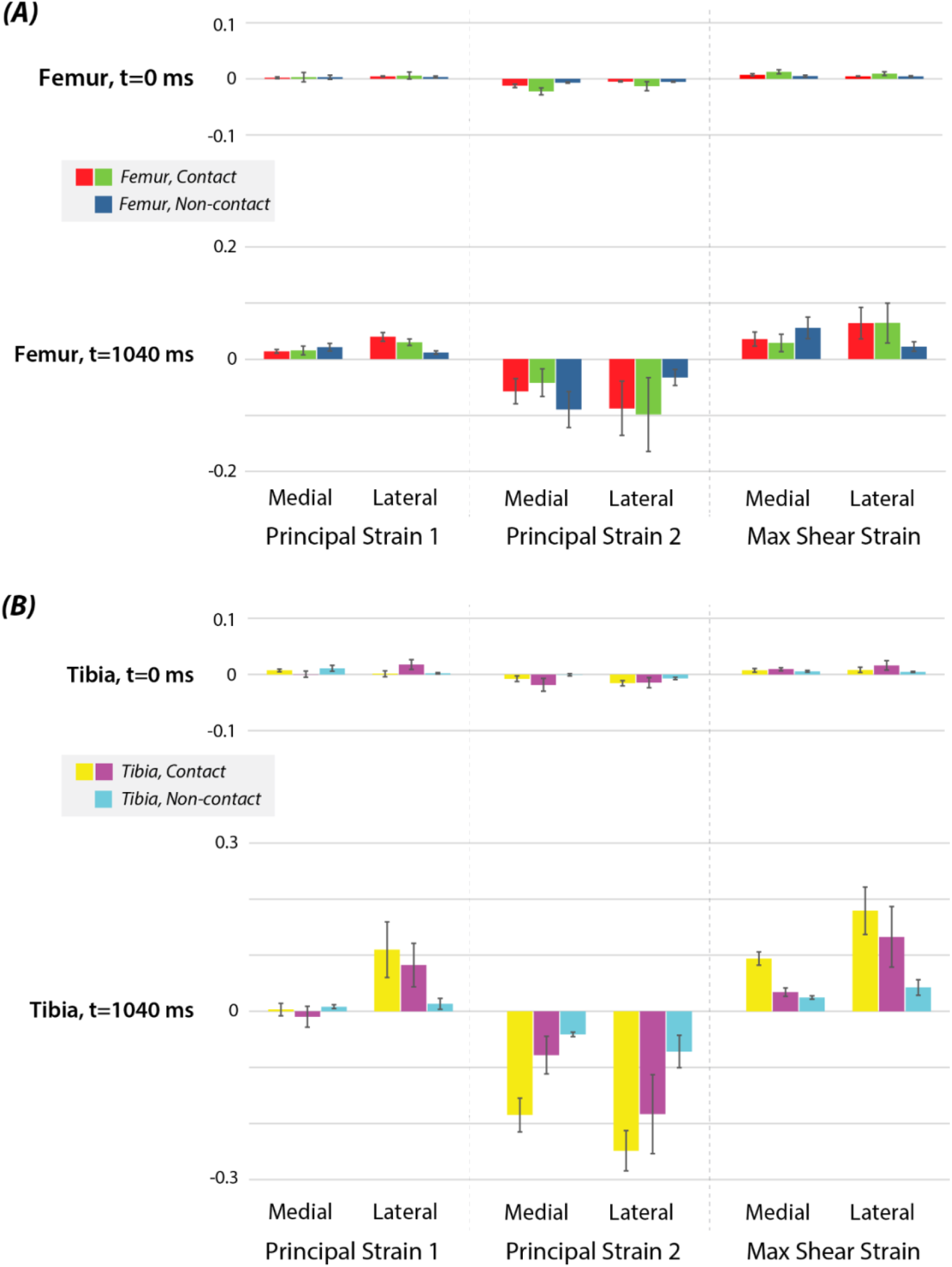
Regional analysis on intact joints showed the increase of principal strains and max shear strain due to load where maximum strain increase was found in the tibia contact regions. Strains in the (A) femur regions and (B) tibia regions increased at t=1040 ms compared to t=0 ms where principal strain 1 and 2 showed a similar trend with E_*xx*_ and E_*yy*_, respectively. Error bars=SEM (N=3).

**Figure S5.**
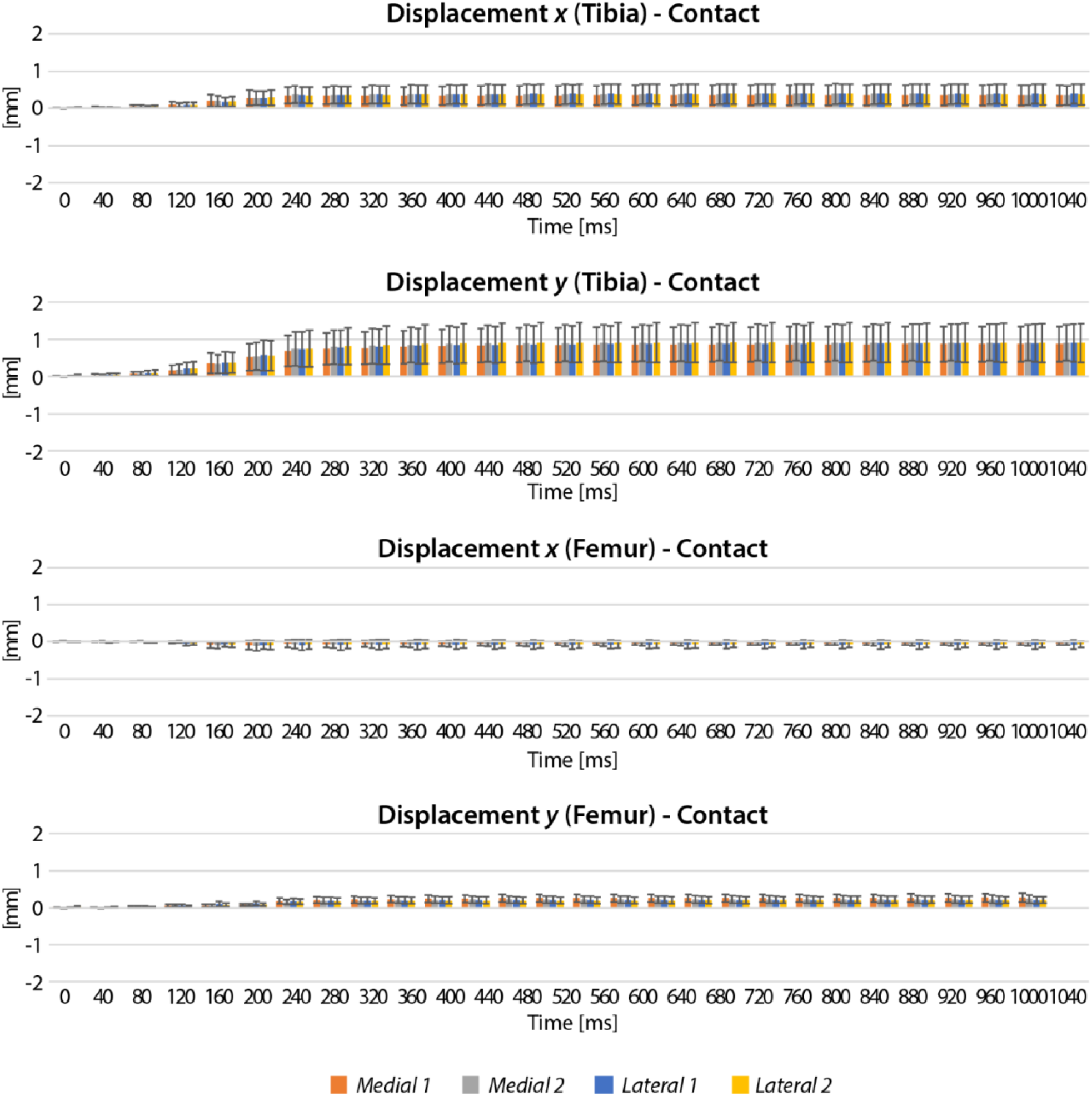
Displacements of intact joints in each time point displayed an increase for contact regions. Contact tibia regions had higher values compared to femur and displacement *y* overall was greater than *x*. Medial 1, lateral 1 refer to the middle regions (2, 5, 8, 11), and medial 2, lateral 2 indicate the inner regions (3, 6, 7, 10) among the contact regions.

**Figure S6.**
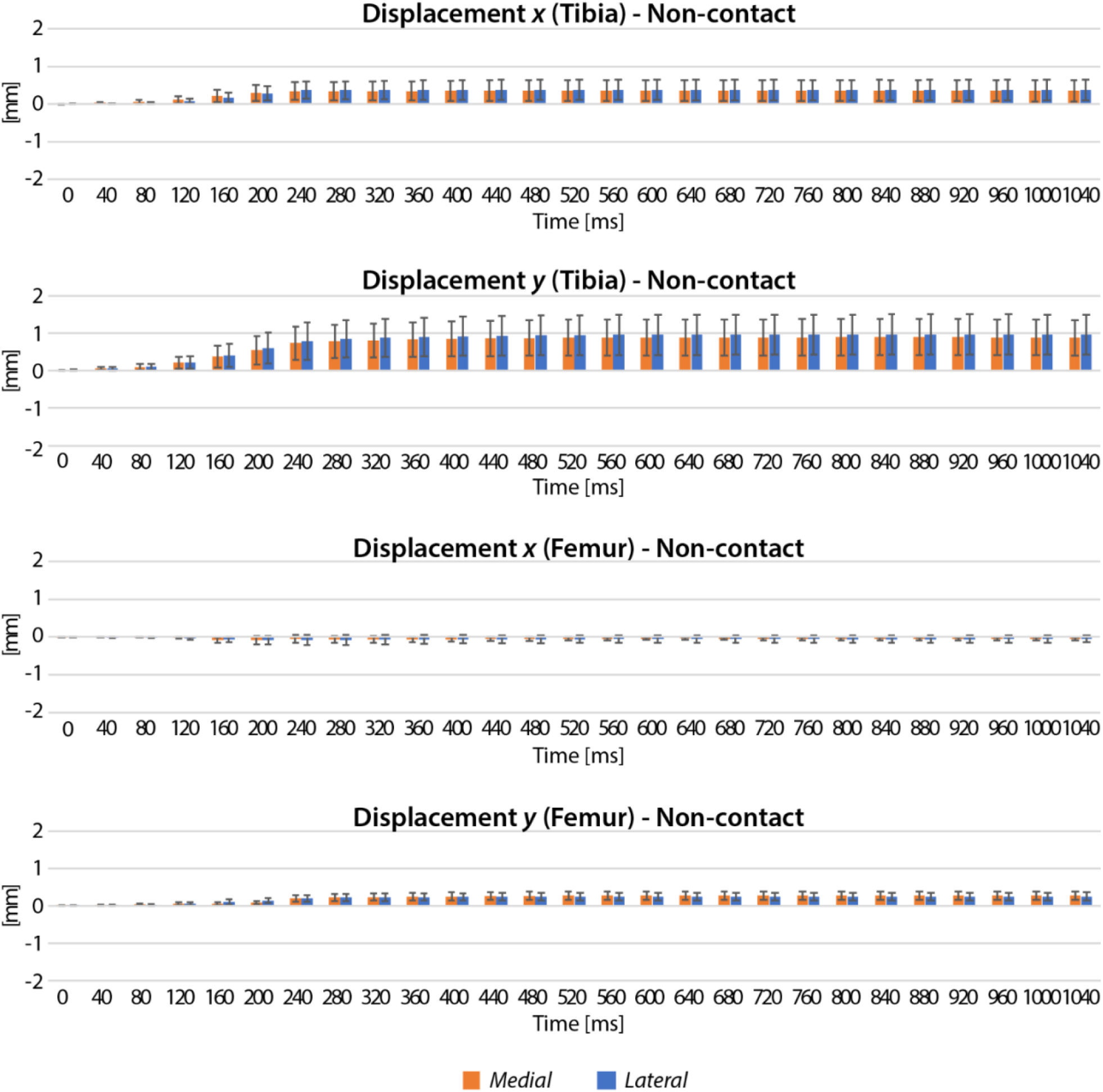
Displacements of intact joints in each time point displayed an increase for non-contact regions. Non-contact regions had a similar trend with the contact regions.

**Figure S7.**
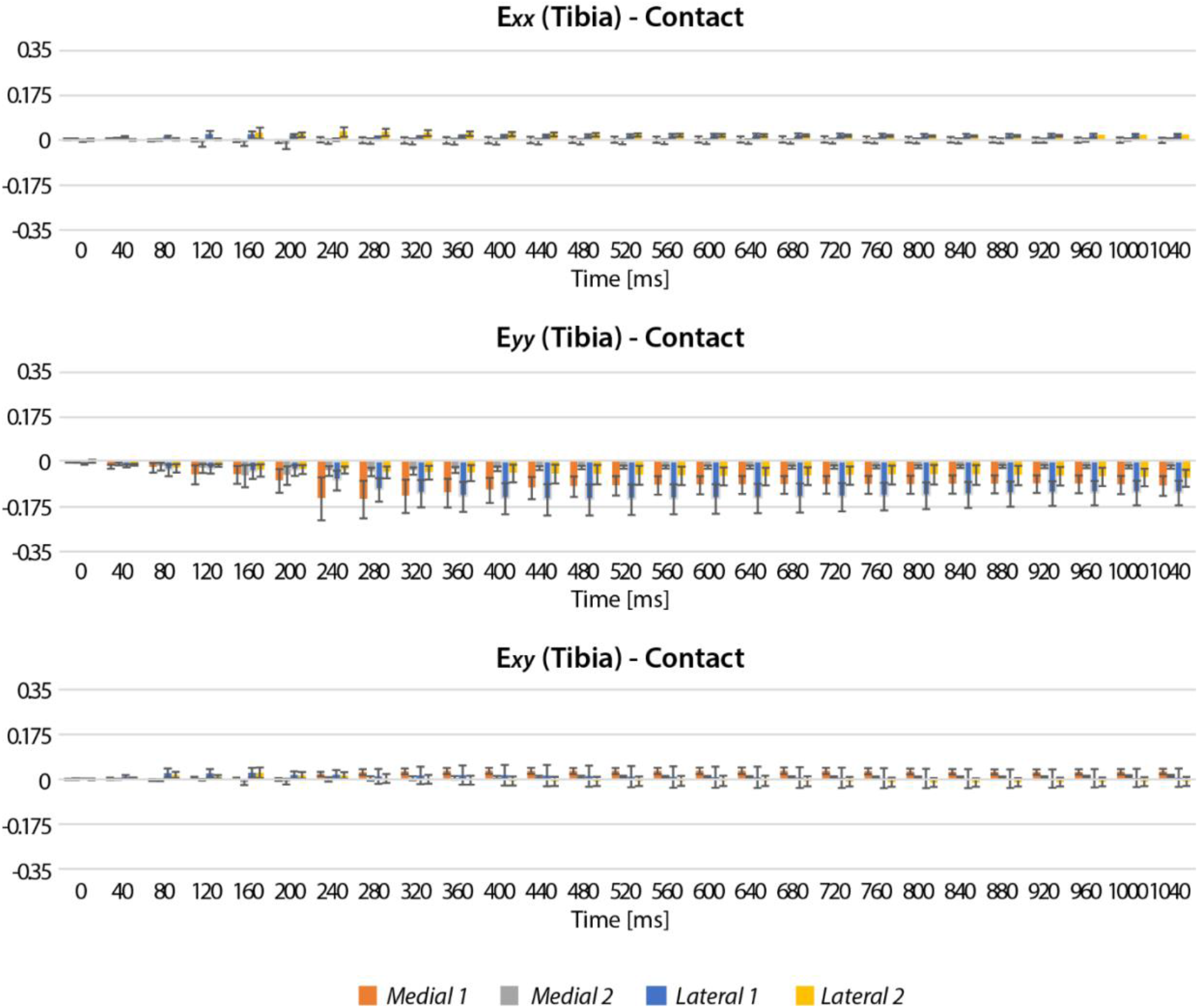
Strains of intact joints in each time point displayed an increase for tibia contact regions. Among the contact regions, E_*yy*_ had maximum magnitudes showing the increase over time. Medial 1, lateral 1 refer to the middle regions (2, 5, 8, 11), and medial 2, lateral 2 indicate the inner regions (3, 6, 7, 10) among the contact regions.

**Figure S8.**
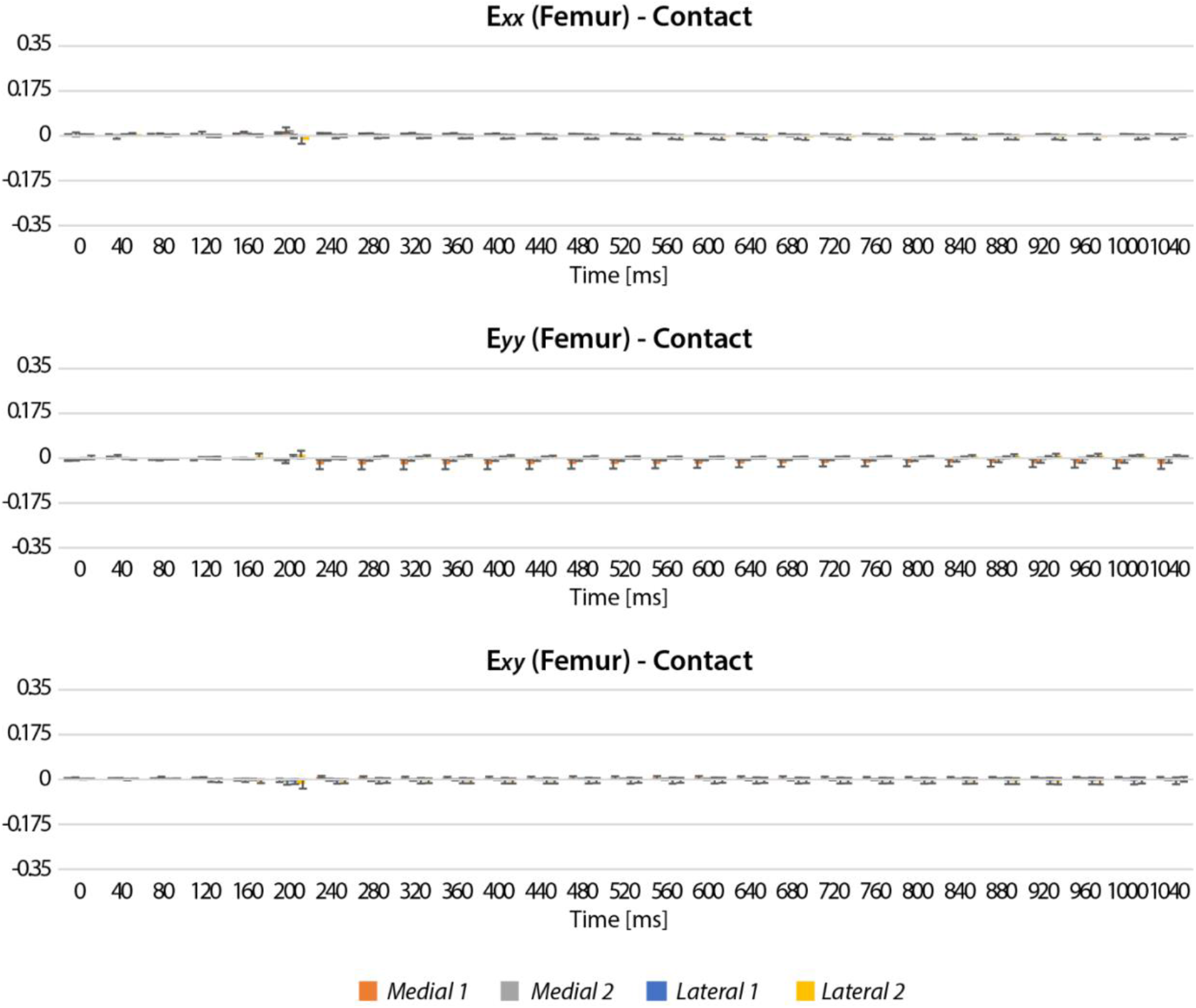
Strains of intact joints in each time point displayed an increase for femur contact regions. Strain values were smaller than the tibia contact regions. Medial 1, lateral 1 refer to the middle regions (2, 5, 8, 11), and medial 2, lateral 2 indicate the inner regions (3, 6, 7, 10) among the contact regions.

**Figure S9.**
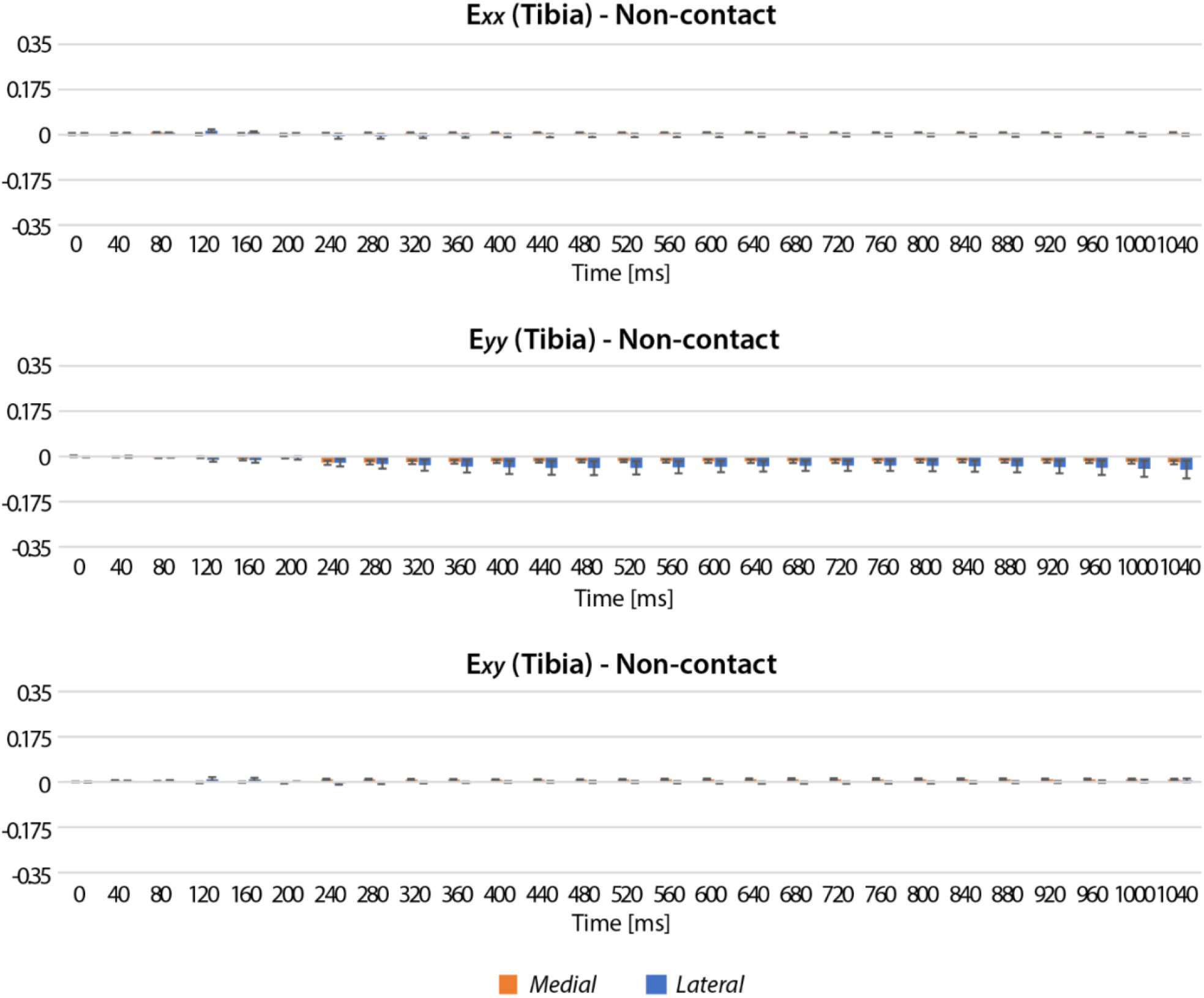
Strains of intact joints in each time point displayed an increase for tibia non-contact regions. Non-contact regions also showed the highest magnitudes in E_*yy*_ however substantially smaller than contact regions.

**Figure S10.**
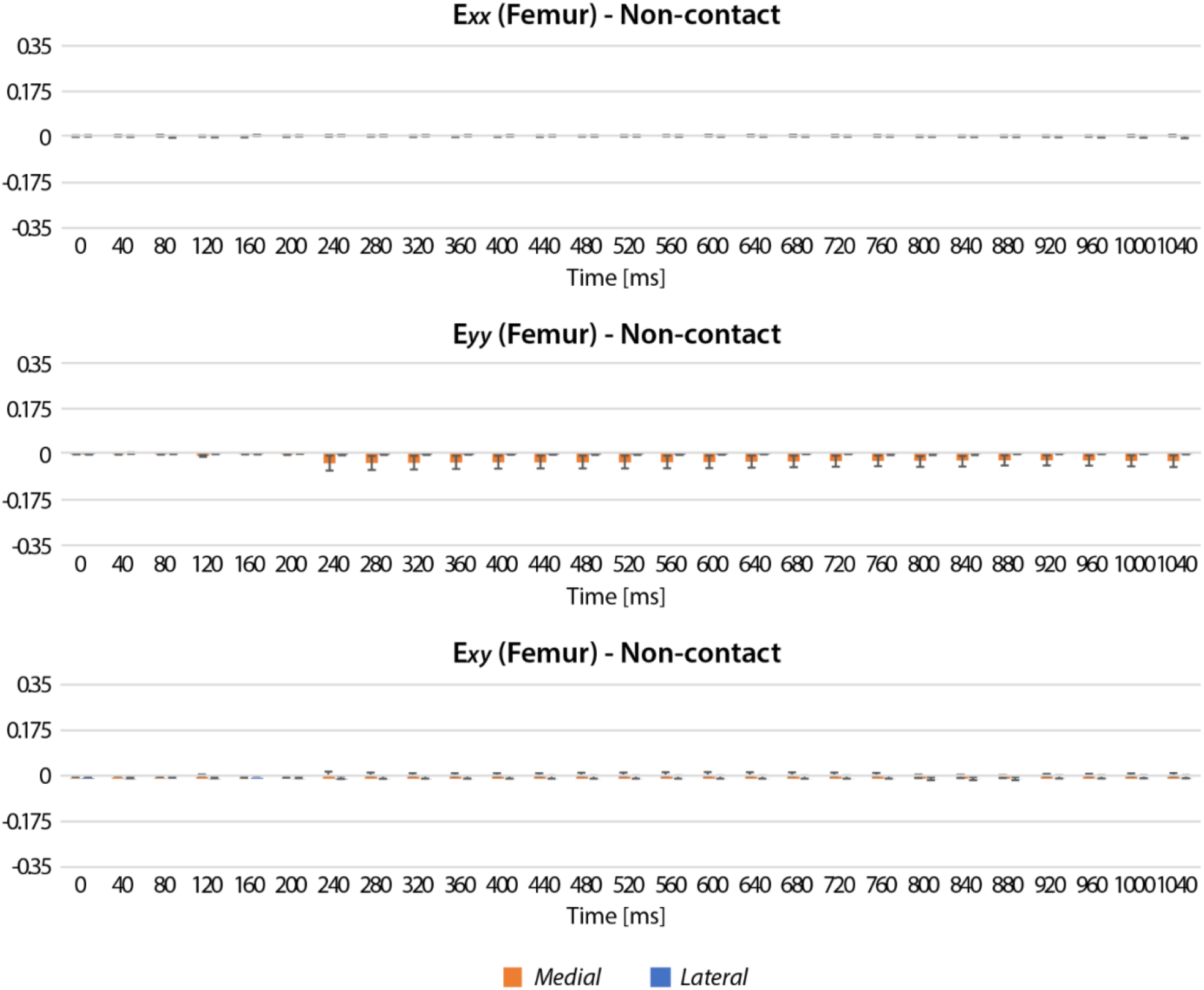
Strains of intact joints in each time point displayed an increase for femur non-contact regions. Femur non-contact regions had a similar trend with the tibia non-contact regions.

**Figure S11.**
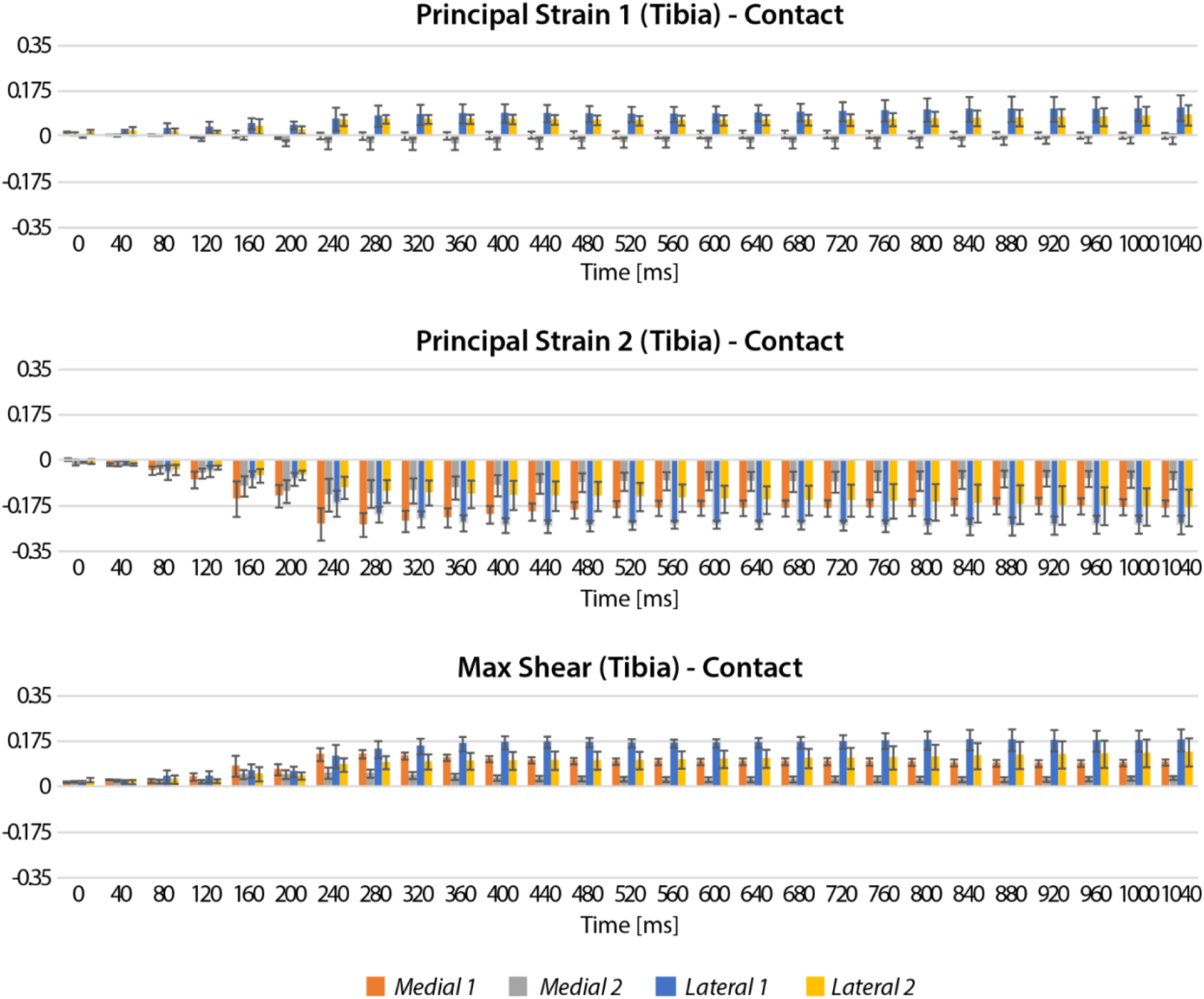
Principal strains and max shear strain of intact joints in each time point displayed an increase for tibia contact regions. In the tibia contact regions, principal strain 2 had the highest magnitudes which increased with time. Medial 1, lateral 1 refer to the middle regions (2, 5, 8, 11), and medial 2, lateral 2 indicate the inner regions (3, 6, 7, 10) among the contact regions.

**Figure S12.**
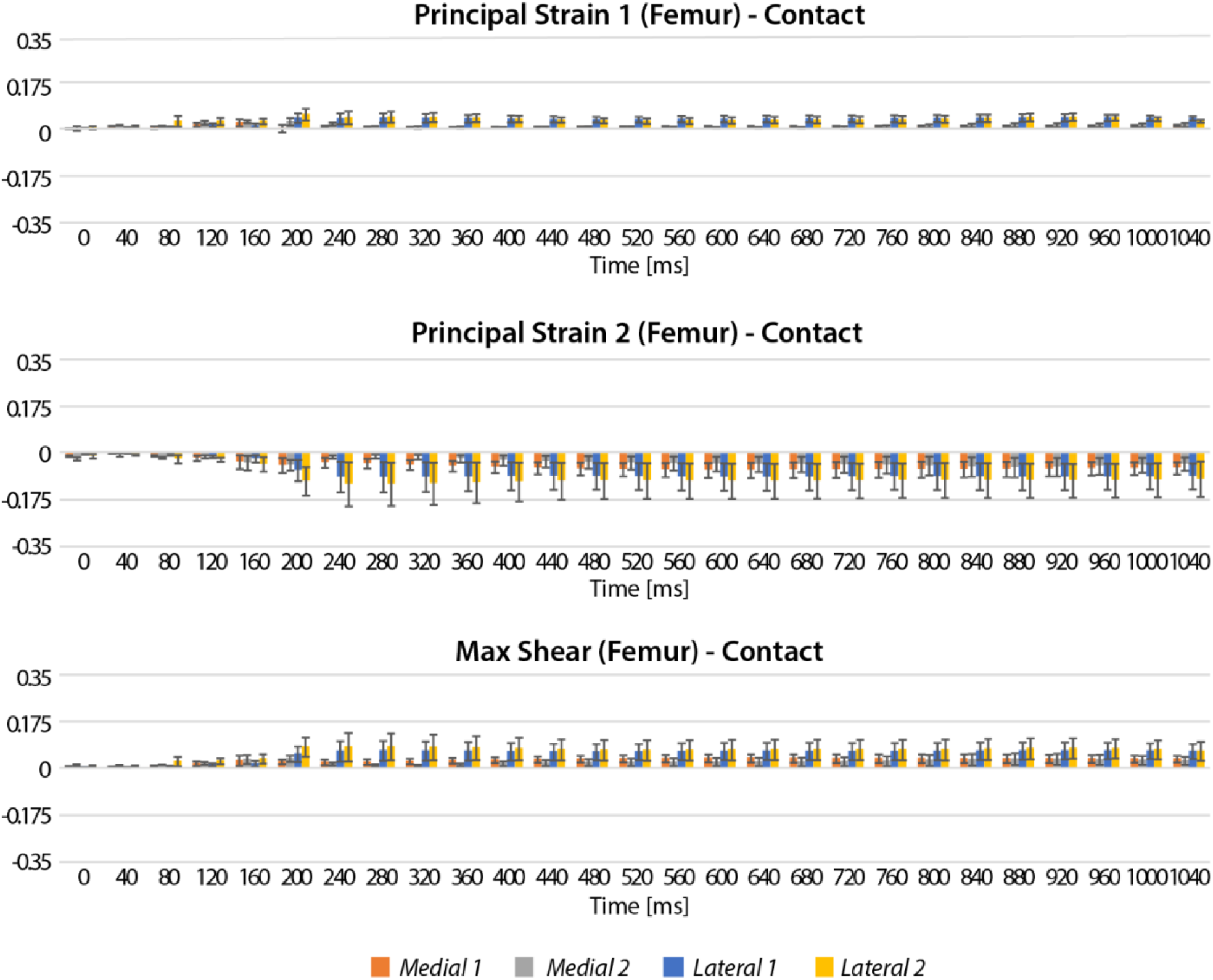
Principal strains and max shear strain of intact joints in each time point displayed an increase for femur contact regions. Similar with the tibia contact regions, the femur contact regions had the highest magnitudes in principal strain 2 which increased with time. Medial 1, lateral 1 refer to the middle regions (2, 5, 8, 11), and medial 2, lateral 2 indicate the inner regions (3, 6, 7, 10) among the contact regions.

**Figure S13.**
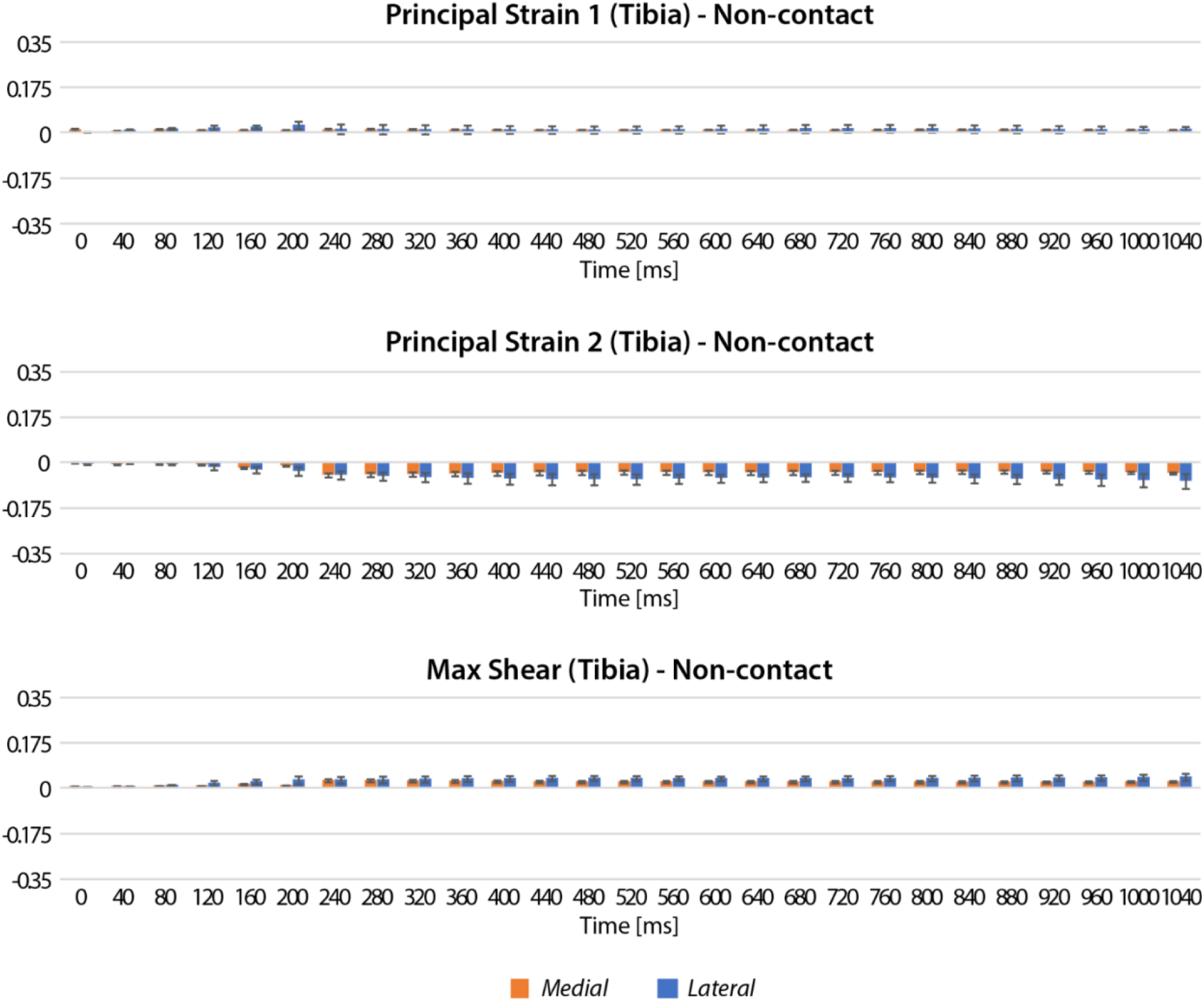
Principal strains and max shear strain of intact joints in each time point displayed an increase for tibia non-contact regions. The tibia non-contact regions showed a similar trend with the contact regions with a substantially smaller magnitude.

**Figure S14.**
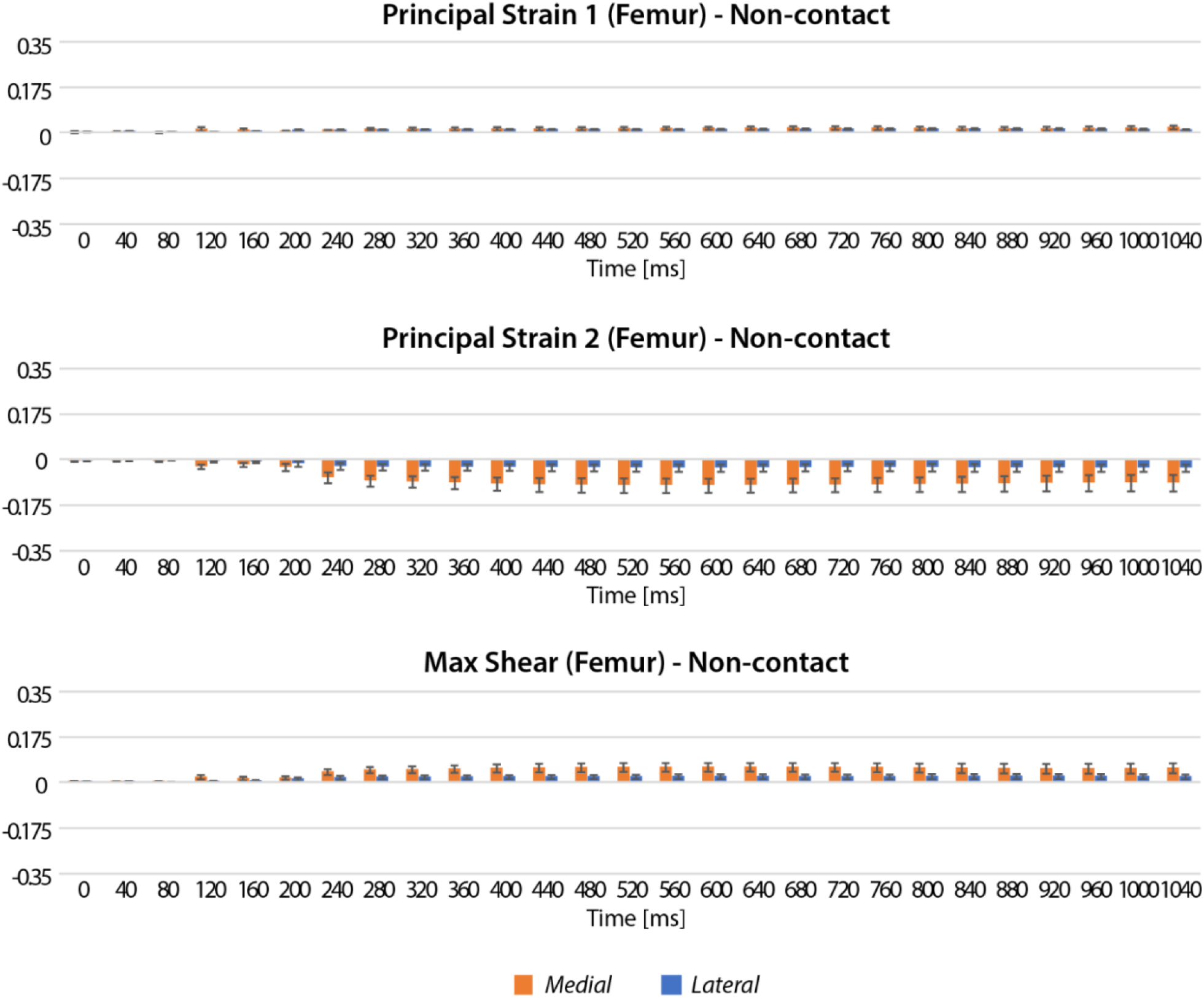
Principal strains and max shear strain of intact joints in each time point displayed an increase for femur non-contact regions. The femur non-contact regions showed a similar trend with the contact regions with a substantially smaller magnitude.

**Figure S15.**
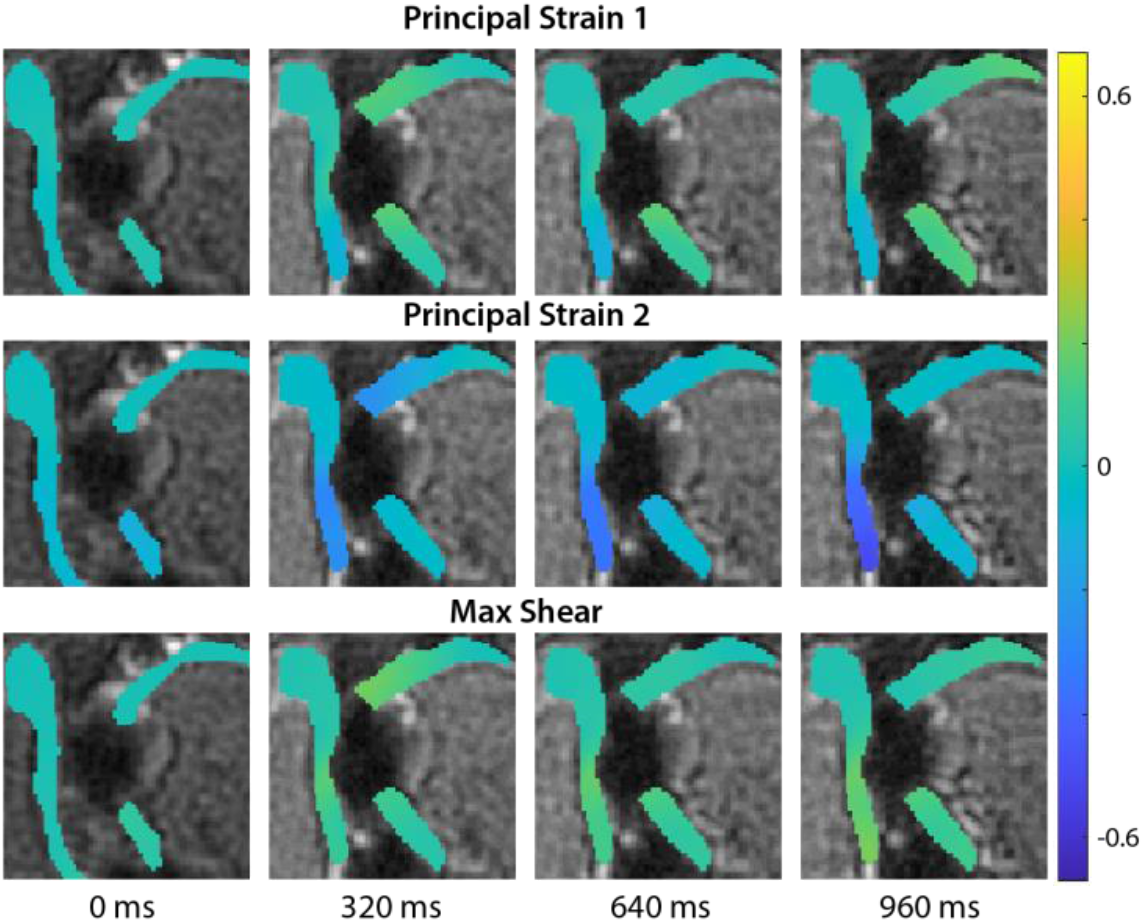
Principal strain 1, 2, and max shear strain obtained by spiral DENSE MRI on a defected bovine joint changed over time. Principal strain 1 and 2 showed a similar trend with E_*xx*_, E_*yy*_ while max shear strain had only positive values as opposed to E_*xy*_ due to the fundamental equation. Analogous to E_*yy*_, higher values of principal strain 2 were observed compared to the intact joint.

**Figure S16.**
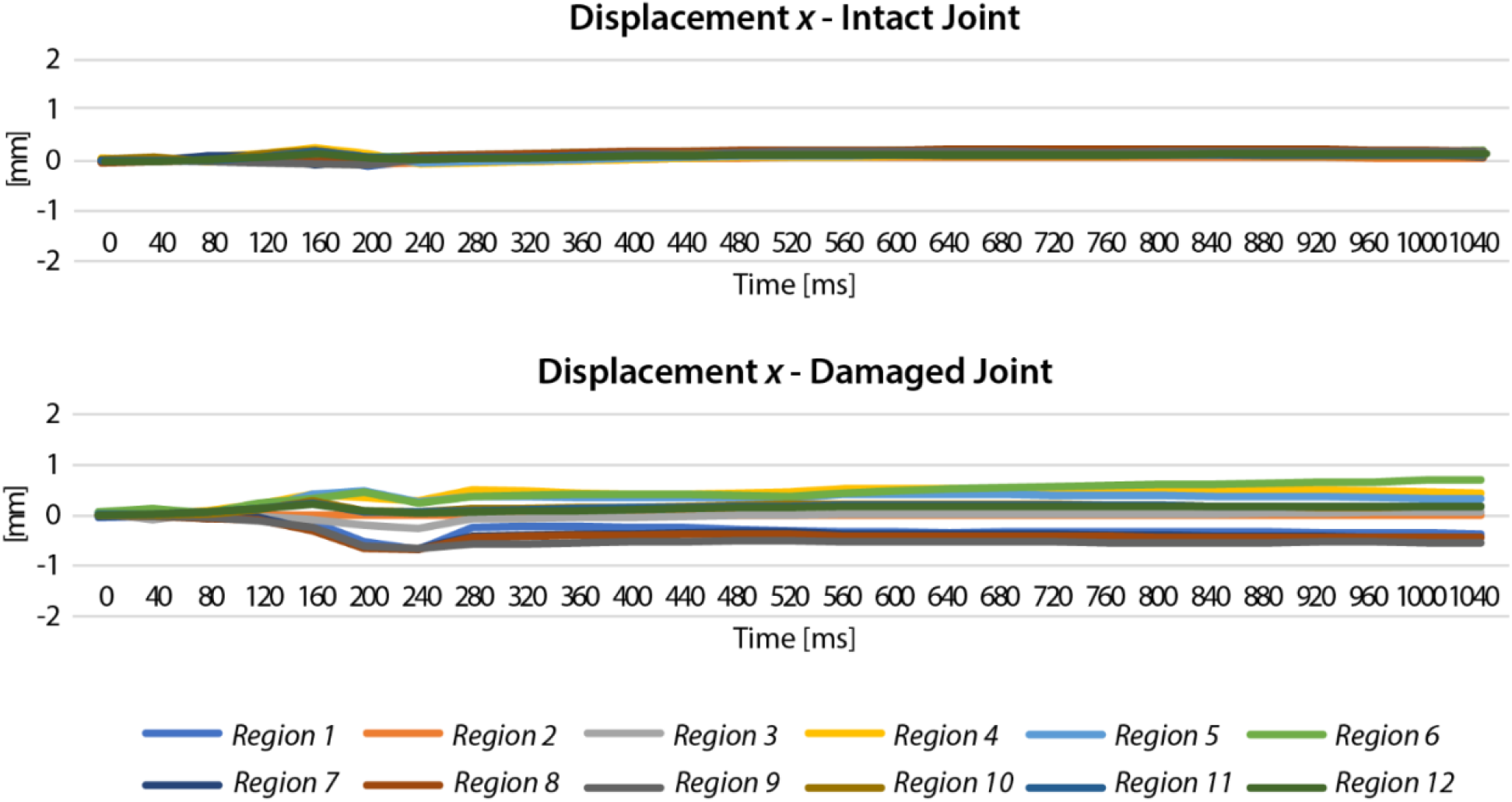
A comparison of intact versus damaged joint in displacement showed damaged joint having greater displacement *x* values. Displacement *x* in the intact sample did not show a significant change over time while the defected sample moved up to 0.7 mm.

**Figure S17.**
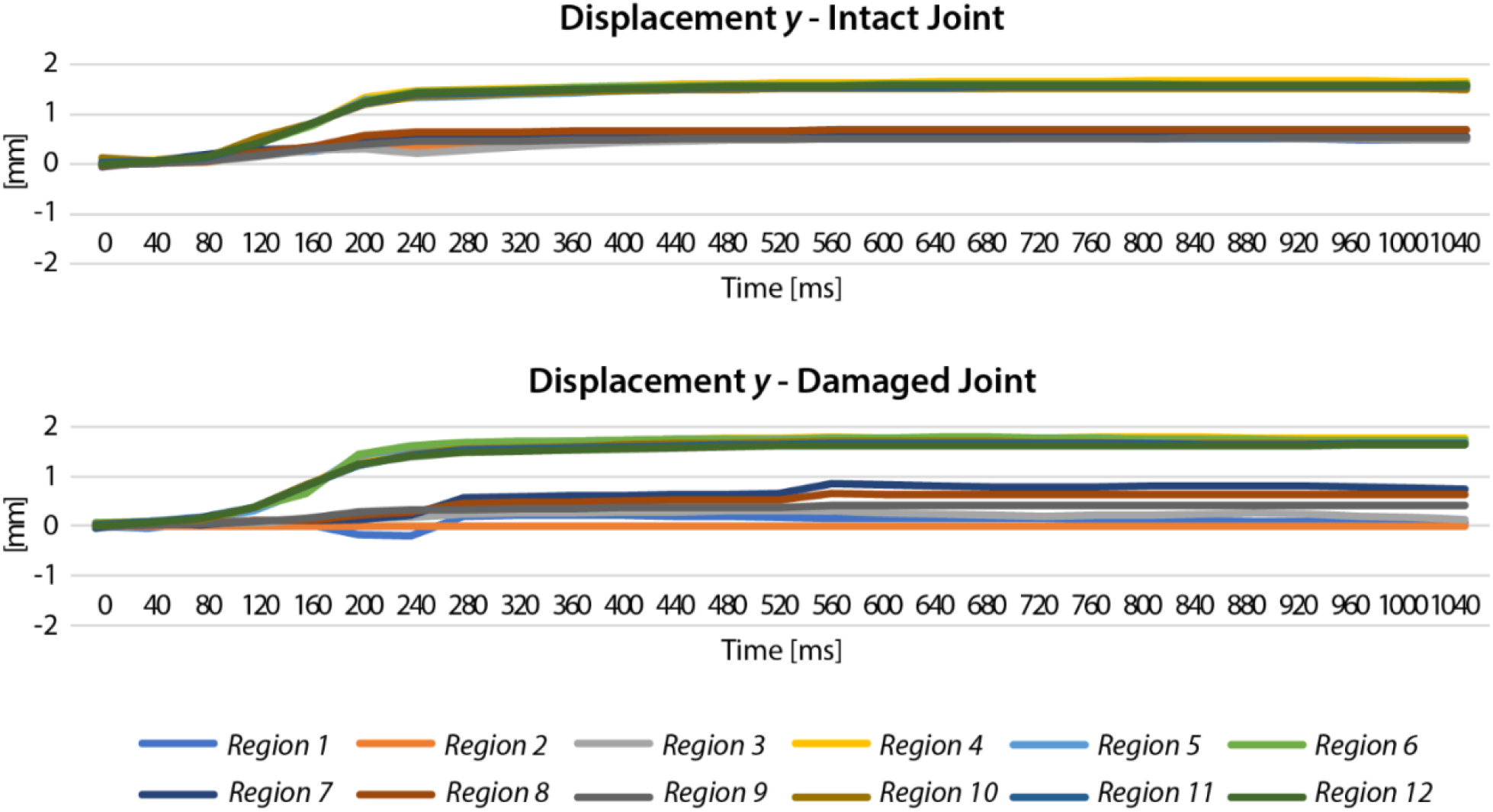
A comparison of intact versus damaged joint in displacement showed damaged joint having greater displacement *y* values. Displacement *y* showed a gradual change over time on both intact and damaged joint where the maximum values were higher in the damaged joint.

**Figure S18.**
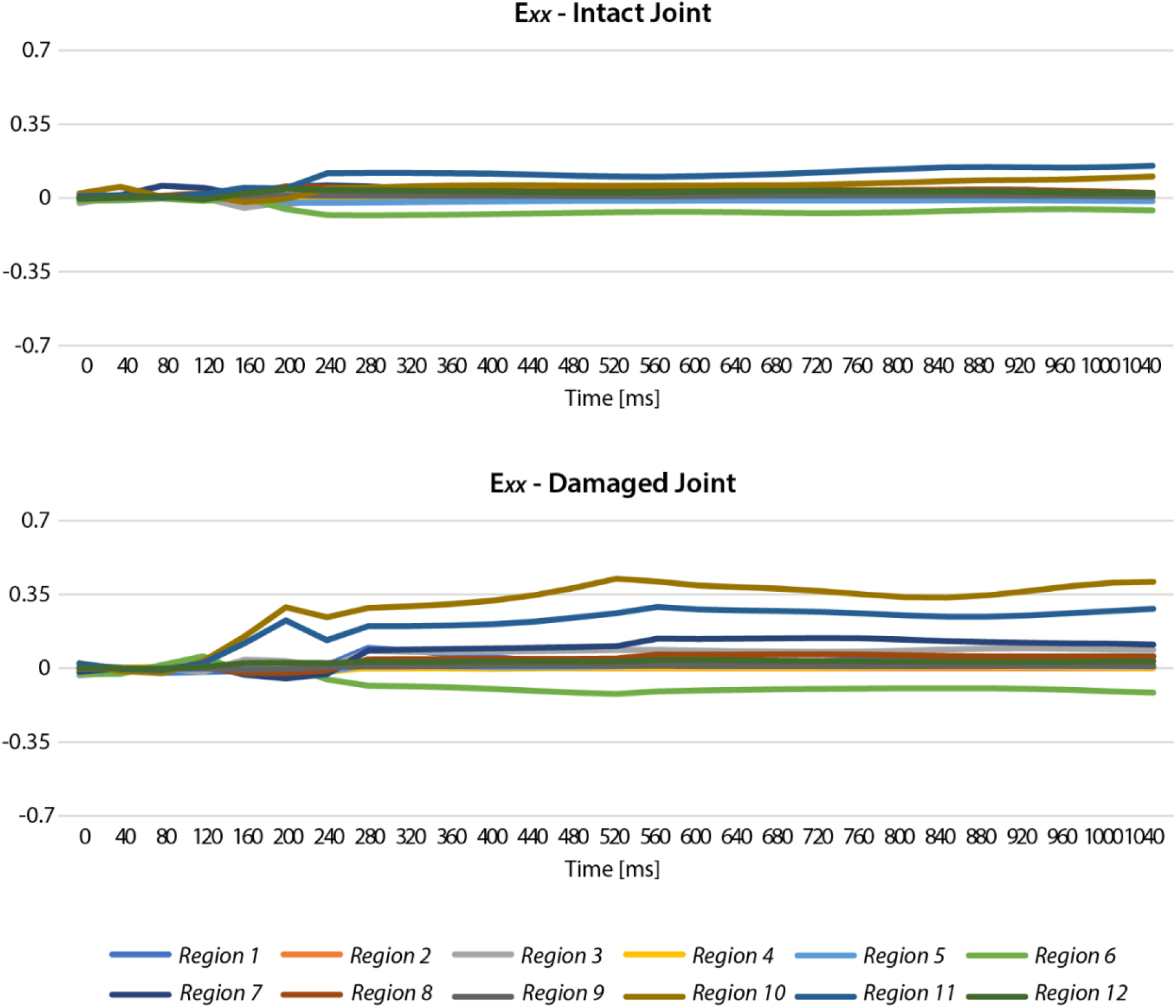
Comparison of intact versus damaged joint in E_*xx*_ showing higher strain values in the damaged joint. E_*xx*_ strain distinctly showed higher magnitudes in the damaged joint mainly in the tibiofemoral contact regions.

**Figure S19.**
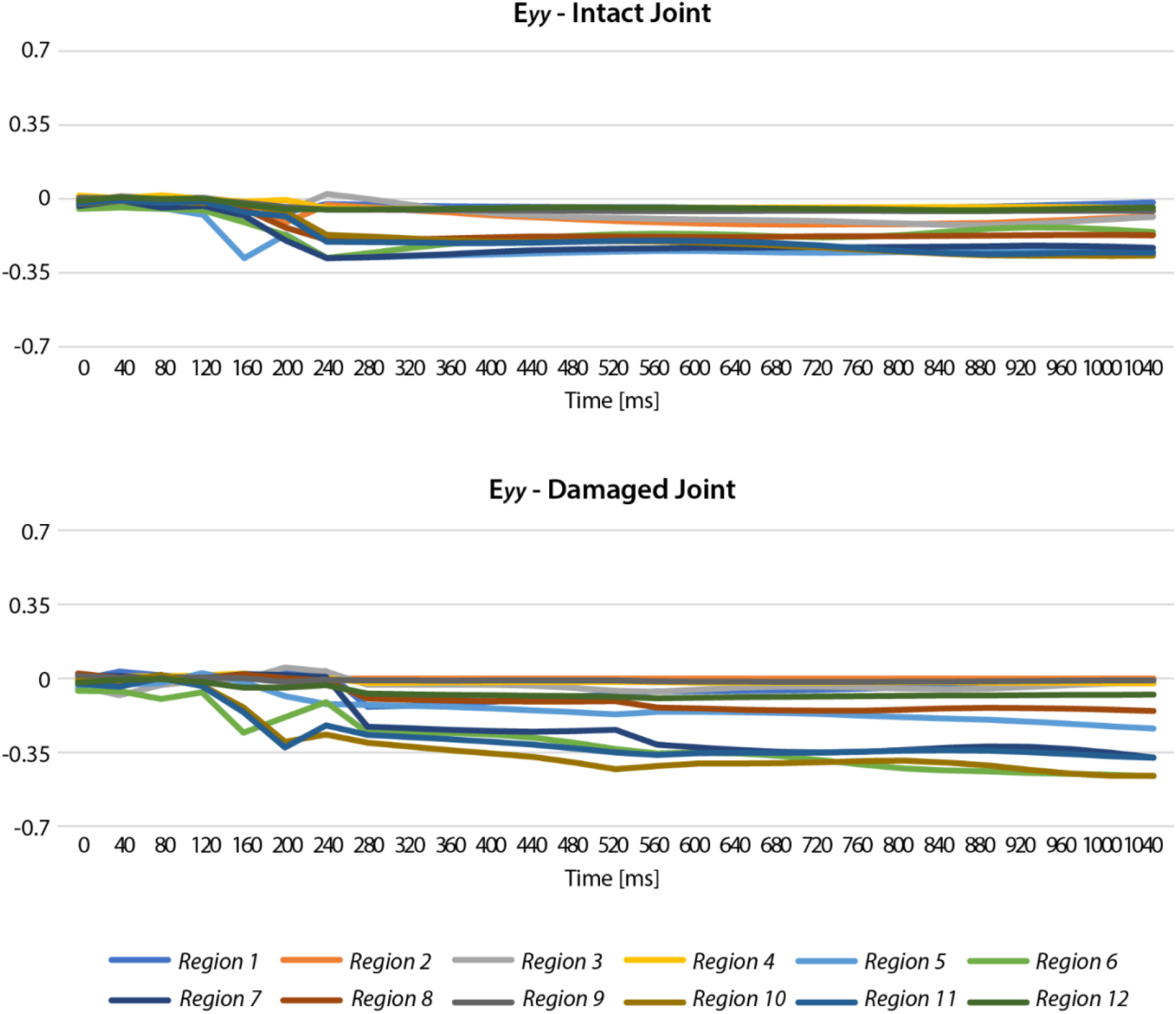
Comparison of intact versus damaged joint in E_*yy*_ showing higher strain values in the damaged joint. E_*yy*_ strain distinctly showed higher magnitudes in the damaged joint mainly in the tibiofemoral contact regions.

**Figure S20.**
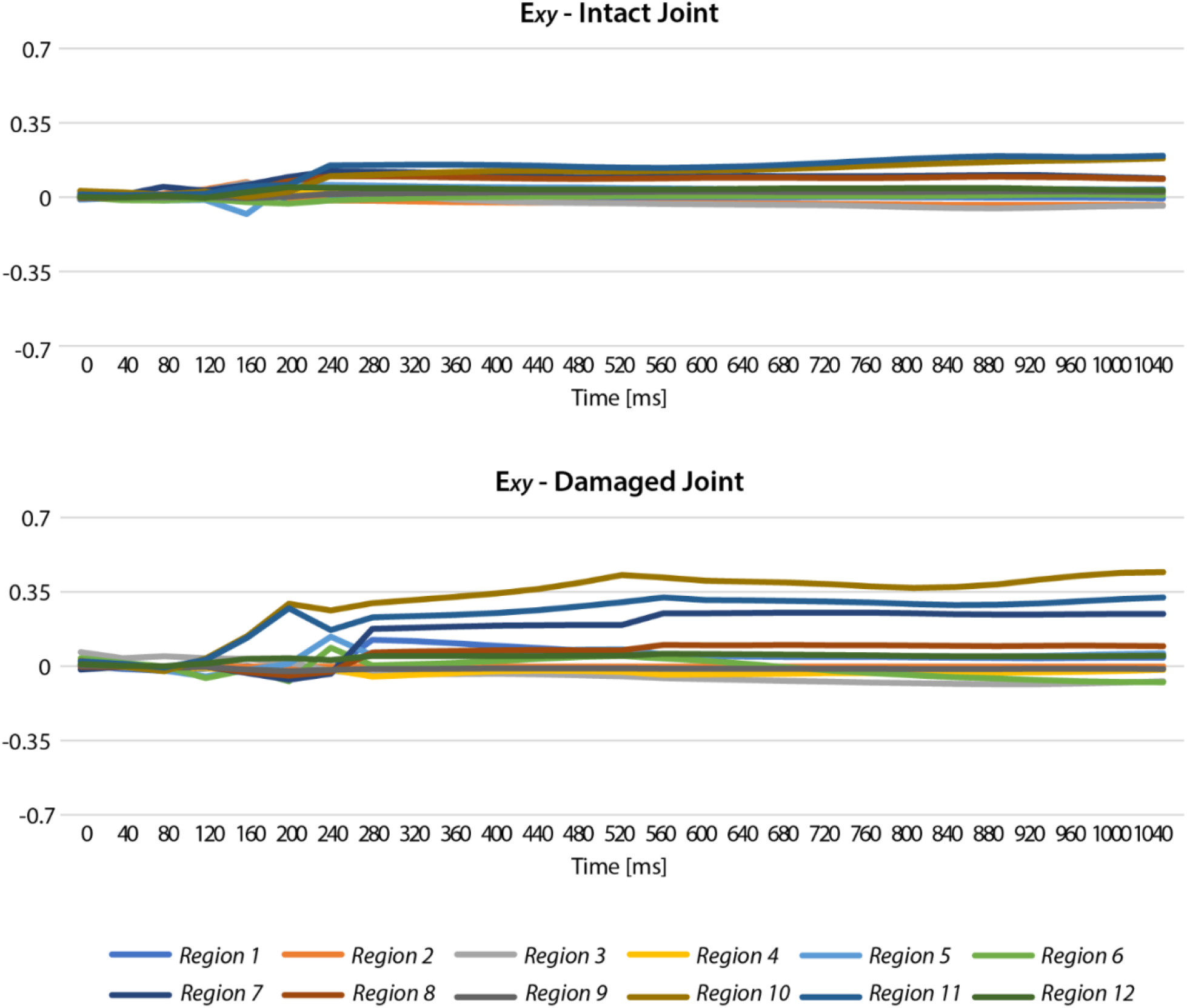
Comparison of intact versus damaged joint in E_*xy*_ showing higher strain values in the damaged joint. E_*xy*_ strain distinctly showed higher magnitudes in the damaged joint mainly in the tibiofemoral contact regions.

**Figure S21.**
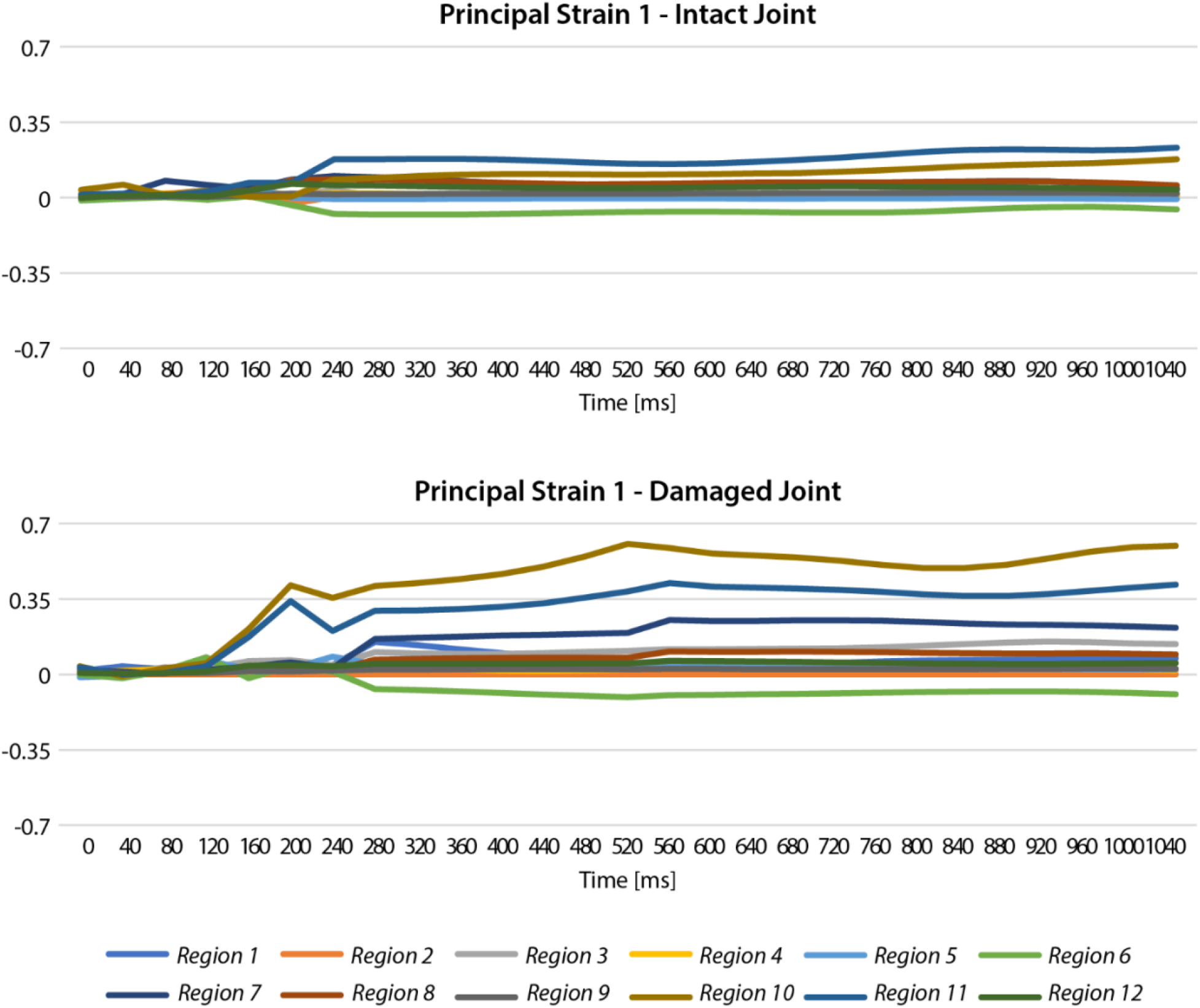
By comparing intact with damaged joint in principal strain 1, damaged joint had higher strain values compared to intact. The magnitude of principal strain 1 increased significantly after defect was generated, mainly in the tibiofemoral contact regions.

**Figure S22.**
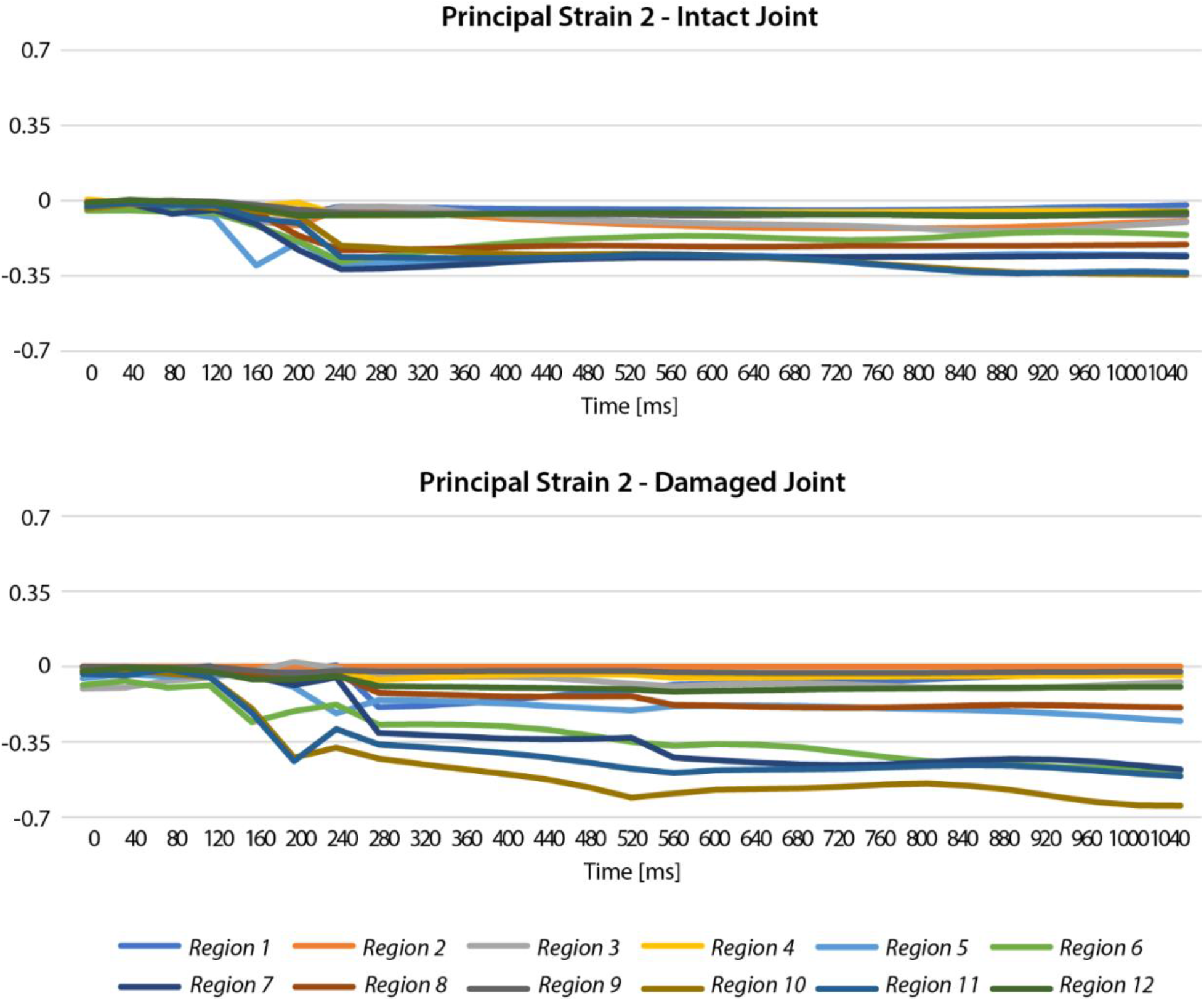
By comparing intact with damaged joint in principal strain 2, damaged joint had higher strain values compared to intact. The magnitude of principal strain 2 increased significantly after defect was generated, mainly in the tibiofemoral contact regions.

**Figure S23.**
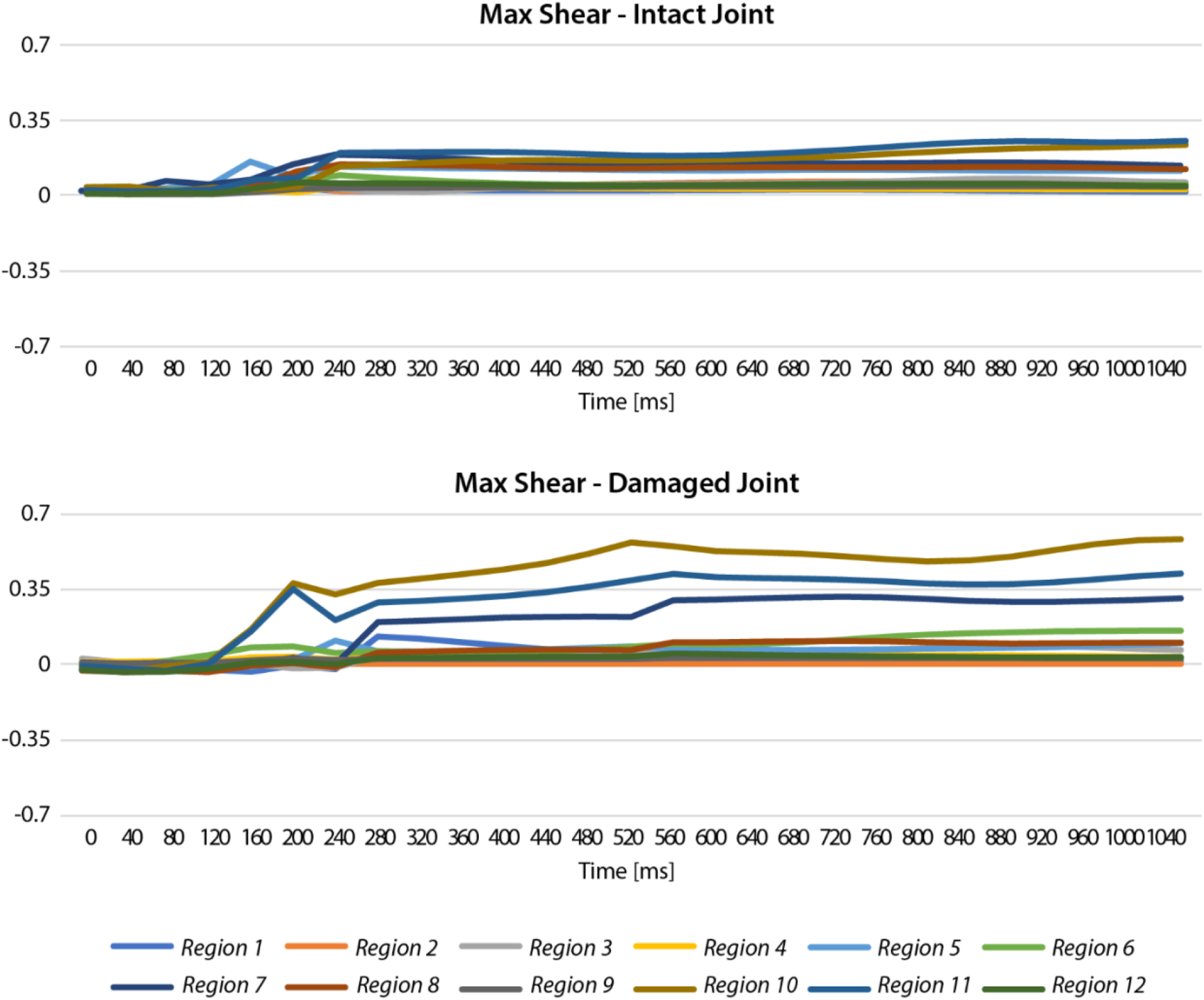
By comparing intact with damaged joint in max shear strain, damaged joint had higher strain values compared to intact. The magnitude of max shear strain increased significantly after defect was generated, mainly in the tibiofemoral contact regions.

